# Single-Cell Multi-Omic Roadmap of Human Fetal Pancreatic Development

**DOI:** 10.1101/2022.02.17.480942

**Authors:** de la O Sean, Zhe Liu, Han Sun, Shengyang K. Yu, Daniel M. Wong, Emily Chu, Sneha A. Rao, Nicolas Eng, Gabriel Peixoto, Jacquelyn Bouza, Yin Shen, Sarah M. Knox, Aaron D. Tward, Anna L. Gloyn, Julie B. Sneddon

## Abstract

The critical cellular transitions that govern human pancreas development are largely unknown. We performed large-scale single-cell RNA-sequencing (scRNA-Seq) to interrogate human fetal pancreas development from 8-20 weeks post conception. We identified 103 distinct cell types, including four novel endocrine progenitor subtypes displaying unique transcriptional features and differentiation potency. Integration with single-nucleus Assay for Transposase Accessible Chromatin Sequencing (snATAC-Seq) identified candidate regulators of human endocrine cell fate and revealed development-specific regulatory annotation at diabetes risk loci. Comparison of *in vitro* stem cell-derived and endogenous endocrine cells predicted aberrant genetic programs leading to the generation of off-target cells. Finally, knock-out studies revealed that the gene *FEV* regulates human endocrine differentiation. This work establishes a roadmap of human pancreatic development, highlights previously unappreciated cellular diversity and lineage dynamics, and provides a blueprint for understanding pancreatic disease and physiology, as well as generating human stem cell-derived islet cells *in vitro* for regenerative medicine purposes.

## INTRODUCTION

Type 1 diabetes (T1D) is a disease of the endocrine pancreas characterized by immune-mediated destruction of insulin-producing beta cells. Beta cell replacement therapy holds great promise for eliminating the need for exogenous insulin delivery and effectively curing the disease (Melton, 2021; Migliorini et al., 2021). Several protocols have been devised to generate insulin-secreting beta-like cells from human pluripotent stem cells (hPSCs) using stepwise differentiation platforms that aim to mimic endogenous development by recapitulating key cell stages through the carefully timed addition and withdrawal of defined combinations of signaling factors (Nostro et al., 2015; Pagliuca et al., 2014; Rezania et al., 2014; Russ et al., 2015; Velazco-Cruz et al., 2020; Veres et al., 2019). These protocols suffer, however, from the production of non-endocrine cell types and a failure to match the transcriptional profiles and glucose responsiveness of primary adult human islets. This may be due to a relative lack of understanding about human endocrine development *in vivo*, as current protocols are based on knowledge of rodent development and may therefore be missing key regulatory pathways and lineage steps unique to human development. Indeed, multiple studies have identified discrepancies between mouse and human pancreatic islets, including structural (Dolenšek et al., 2015), transcriptomic (Baron et al., 2016), and metabolic (MacDonald et al., 2011) differences. Therefore, gaining a deeper understanding of human endocrine development is crucial for continued progress towards generating *in vitro*-derived beta-like cells that recapitulate endogenous function.

In mice, all pancreatic epithelial cell types, including both exocrine and endocrine, are generated from a domain of the gut tube that begins to express the transcription factor Pdx1 around E8.5 (Ohlsson et al., 1993). The initial pancreatic bud then branches extensively, forming the tip and trunk regions of the finger-like projections that comprise the ductal epithelium as it expands and begins regional specification (Shih et al., 2013). The hormone-expressing endocrine cells, including insulin-producing beta cells, derive from endocrine progenitor (EP) cells that activate the expression of the transcription factor Neurog3 in a subset of ductal trunk cells (Salisbury et al., 2014). During human development, *NEUROG3* is also a presumed marker of EP cells (Salisbury et al., 2014). The expression of NEUROG3 in humans begins as early as 8 weeks post conception (w), peaks at 10-12 w, and then gradually decreases to very low levels by 35 w (Salisbury et al., 2014). While in murine pancreatic development the expression of Neurog3 in EPs occurs in two distinct waves, in human development it has been reported to occur in a single wave, further highlighting differences between human and rodent pancreas development (Villasenor et al. 2008; Jennings et al. 2015).

Heterotypic interactions between epithelial and non-epithelial cells are broadly important for mammalian development, including in the pancreas, where they regulate expansion of pancreatic epithelial progenitors as well as their subsequent differentiation (Bhushan et al., 2001; Cleaver and Dor, 2012; Golosow and Grobstein, 1962; Landsman et al., 2011). In particular, islet development depends on complex interactions between endocrine cells and multiple other cell types, including neurons and endothelial cells (Borden et al., 2013; Cleaver and Dor, 2012; Lammert et al., 2001). To fully understand human endocrine pancreas development *in vivo* and then successfully mimic the process *in vitro* will require a comprehensive catalogue of not only the endocrine cells, but also the non-epithelial pancreatic cell types, as well as signaling pathways through which they act.

Murine single-cell RNA-sequencing (scRNA-Seq) studies performed by our laboratory and others have uncovered EP subtypes downstream of the *Neurog3*-expressing population, and some have catalogued the cell heterogeneity within other non-endocrine compartments as well (Bastidas-Ponce et al., 2019; Byrnes et al., 2018; Yu et al., 2019b). Single-cell studies of human pancreas have begun to reveal heterogeneity of human endocrine cells. For instance, scRNA-Seq has been applied to cells generated by beta cell differentiation protocols *in vitro* and to adult human islets, where previously unappreciated levels of cellular heterogeneity were described (Baron et al., 2016; Gonçalves et al., 2021; Muraro et al., 2016; Petersen et al., 2017; Segerstolpe et al., 2016; Veres et al., 2019; Xin et al., 2018; Yu et al., 2021). A study of human pancreas tissue at 9 w used single-cell qPCR on a small number of sorted cells to detect the expression of 96 prospectively defined developmental genes and described a putative EP population as well as an early differentiated endocrine cell population (Ramond et al., 2018). More recently, a paper focusing on early pancreatic epithelial progenitors reported potential pathways that may be broadly mediating interactions between mesenchymal and epithelial cells; however, the number of endocrine cells investigated was limited (Gonçalves et al., 2021). Work focusing on human fetal endocrinogenesis identified putative EP cell states *in silico*, although these states remain to be confirmed *in vivo* and there was no investigation of how endocrine cells interact with other cell types in the developing pancreas (Yu et al., 2021). Thus, a comprehensive characterization of the full panoply of both epithelial and non-epithelial cell types in the developing human pancreas is still lacking. Importantly, the field also still lacks an understanding at the single-cell level of how cell composition and lineage trajectories of *in vitro* stem cell-derived beta cells compare to those of endogenous developing human cells.

In the quest to characterize the relevant cell states of endocrine differentiation *in vivo* that need to be recapitulated *in vitro* for cell replacement therapy approaches, the field also lacks an understanding of the epigenetic mechanisms by which those endocrine cell states are established and maintained. A study investigating *in vitro* generation of pancreatic progenitors identified heterogeneity in global chromatin accessibility depending on which *in vitro* differentiation protocol was used, highlighting the need for an *in vivo* comparator against which epigenetic data from *in vitro* differentiation platforms can be benchmarked (Wesolowska-Andersen et al., 2020). Recent studies using single-nucleus Assay for Transposase-Accessible Chromatin with Sequencing (snATAC-Seq) have provided evidence of distinct epigenetic states across endocrine subclusters in the adult islet (Chiou et al., 2021a). These studies also localized type 1 and type 2 diabetes-associated genetic risk variants to regions of accessible chromatin in adult islet cells and predicted their regulatory function by interpreting their co-accessibility with target genes (Chiou et al., 2021b, 2021a; Rai et al., 2020). Although a large number of diabetes genetic risk variants have been discovered in Genome-Wide Association Studies (GWAS) (Mahajan et al., 2018; Pociot, 2017; Robertson et al., 2021), it is not clear which are operant during development, thereby exhibiting regulatory functions in a cell-specific and/or developmental stage-specific manner.

In this study, we utilize large-scale scRNA-Seq to generate a comprehensive atlas of human fetal pancreas tissue ranging from 8 to 20 w. We describe previously unappreciated levels of cell heterogeneity within the endothelial, mesenchymal, exocrine, neuronal, immune, and endocrine lineages of the human pancreas and identify putative signaling pathways active among these lineages. Within the endocrine lineage, we identify four novel progenitor cell types, confirm their existence in independent tissue samples, and reconstruct their lineage trajectories *in silico*. By performing snATAC-Seq analysis on 12 w human fetal pancreatic tissue and performing a multi-omic analysis integrating snATAC-Seq and scRNA-Seq data, we provide novel insights into regulatory landscapes of single cells in the developing human endocrine pancreas. We also leverage the snATAC-Seq data to identify a potential developmental-specific role of multiple diabetes GWAS risk alleles. In addition, we compare the molecular profiles and cellular trajectories of endogenous human fetal pancreatic endocrine cells to those generated *in vitro* from human stem cells. Through genome editing of hPSCs, we identify the transcription factor *FEV* as a regulator of human endocrine differentiation. This study will serve broadly as a resource for the field of pancreatic development and will also provide the foundation for future improvements in therapeutic strategies for generating replacement beta cells that more closely resemble their *in vivo* counterparts both in identity and function.

## RESULTS

### Interrogating the Developing Human Pancreas at Single-Cell Resolution

To characterize the cellular heterogeneity within the human fetal pancreas, we performed droplet-based scRNA-Seq on 8 independent biological samples ranging from 8 to 20 w, a window of human pancreatic development encompassing specification of endocrine progenitors (EPs), differentiation into hormone-producing cell types, and islet morphogenesis (Jennings et al., 2015). Each tissue sample was dissociated and subjected to red blood cell lysis to deplete erythrocytes. Resulting single-cell suspensions were then used directly (“Total Pancreas”) or subjected to magnetic bead-based selection to either enrich for EPCAM+ cells (“Epithelial(+)”), or deplete CD45+ cells (“Immune(-)”), and then prepared for sequencing using the 10x Chromium Single-Cell Gene Expression v3 platform (**Figure 1A**). Following sequencing and quality control processing (see STAR methods), the resulting data were computationally merged into a final dataset consisting of 114,873 cells, each of which was then classified as belonging to a “Broad Group’’ based on the expression of established marker genes: Mesenchymal (*COL3A1*), Immune (*RAC2*), Exocrine (*CPA1*), Endothelial (*PECAM1*), Neuronal (*SOX10*), Endocrine (*CHGA*), and Proliferating (*TOP2A*) (**Figure 1B**). Individual biological samples showed varying degrees of contribution to all Broad Groups, with experimental enrichment or depletion affecting the overall contribution as expected. For instance, samples subjected to positive selection for epithelial cells showed enrichment of endocrine and exocrine populations and relatively fewer cells classified as mesenchymal, immune, or endothelial **(Figure S1A-B)**. Importantly, we observed high concordance of technical and biological replicates based on Pearson correlation of Epithelial cells (**Figure S1C**).

**Figure 1:**
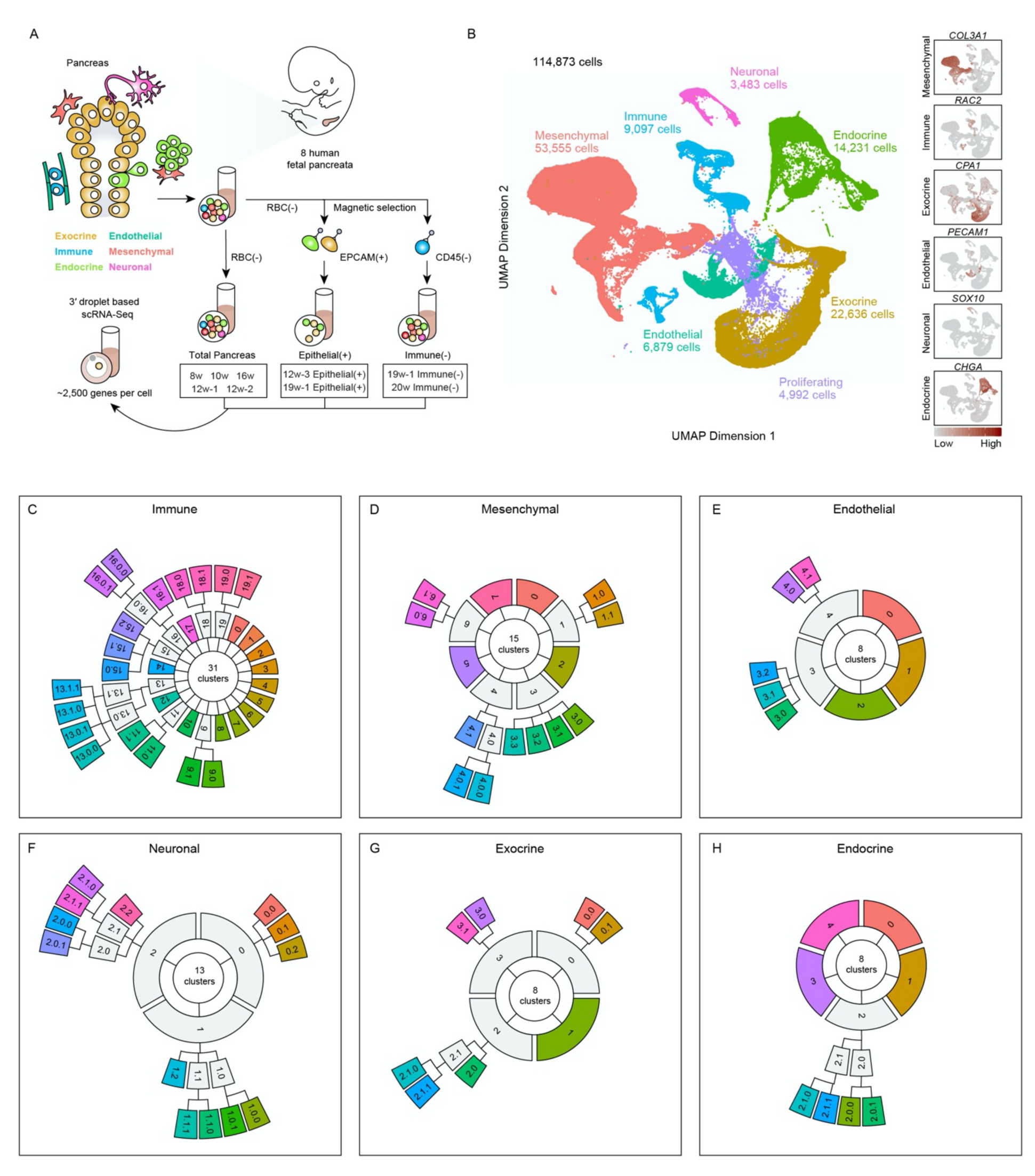
Large-scale single-cell RNA-Sequencing identifies striking cellular heterogeneity within the human fetal pancreas. (A) Overview of experimental approach. Eight samples of human fetal pancreas tissue ranging from 8 to 20 weeks post-conception (w) were dissociated and subjected to red blood cell lysis (“RBC(-)”) to deplete erythrocytes. Resulting single-cell suspensions were then used directly (“Total Pancreas”), or subjected to either magnetic bead-based enrichment for EPCAM+ cells (“Epithelial(+)”), or depletion of CD45+ cells (“Immune(-)”), followed by single-cell RNA-Sequencing (scRNA-Seq). The 19 w-Epithelial(+) and 19 w-Immune(-) cells were from the same tissue sample, and the 12w-1, 12w-2 and 12w-3-Epithelial(+) samples are three independent biological replicates. (B) UMAP visualization of the merged scRNA-Seq dataset from all nine conditions, derived from eight biological specimens at six developmental timepoints. Each cell is color-coded according to the Broad Group to which it belongs. Expression of marker genes *COL3A1, RAC2, CPA1, PECAM1, SOX10,* and *CHGA* are displayed in feature plots to the right. (C)-(H) Iterative clustering performed using the CellFindR algorithm revealed three layers of heterogeneity within each Broad Group, with Tier 1, 2, and 3 populations arranged in the inner, middle, and outer circles, respectively. Clustering hierarchies depict cellular populations within the Broad Groups of (C) immune; (D) mesenchymal; (E) endothelial; (F) neuronal; (G) exocrine; and (H) endocrine cells. All 103 terminal Tier 3 populations identified by CellFindR are colored to match the UMAPs in Figure 2 and Figure S2.

To further investigate the cellular heterogeneity within each Broad Group, we next applied the clustering algorithm CellFindR (Rust et al., 2020; Yu et al., 2019a) to our dataset (see STAR Methods). CellFindR iteratively increases Louvain clustering resolution based on the condition that each cluster expresses a minimum of 10 genes with greater than 2-fold expression in comparison to all other clusters. Once this condition is broken, principal component analysis (PCAs) are recalculated and each cluster is further sub-clustered following the same condition, creating tiers of clusters with the nomenclature of the initial cluster represented by an integer and subsequent sub-clusters followed by a period and an integer. CellFindR defines cell populations that are biologically relevant and generates a clustering map with multi-tier hierarchy. Within the merged datasets, CellFindR identified a total of 103 clusters in the developing human pancreas, including 15 mesenchymal, 8 exocrine, 8 endocrine, 8 endothelial, 13 neuronal, 31 immune, and 20 proliferating clusters. This highlights the striking and previously undescribed cell heterogeneity in the developing human pancreas **(Figure 1C-1H).**

### Atlas of Cellular Heterogeneity and Cell-Cell Communication in Non-Endocrine Lineages of the Fetal Pancreas

The importance of mesenchyme in guiding pancreatic organogenesis has been demonstrated through mechanical removal and recombination experiments, as well as genetic ablation studies (Attali et al., 2007; Golosow and Grobstein, 1962; Landsman et al., 2011). Recent work has shed light on the functional heterogeneity within murine pancreatic mesenchyme, where an Nkx2.5+ mesenchymal sub-population has been reported to establish a pro-endocrine niche during pancreatic development (Cozzitorto et al., 2020). The full panoply of cell subtypes within the human fetal pancreatic mesenchyme and their roles in heterotypic cellular signaling, however, remain unknown. In this study, we employed CellFindR to identify 15 sub-clusters of mesenchymal cells (**Figure 1D**, **Figure 2A**), including known cell types such as vascular smooth muscle (Mature and Immature VSM, clusters 6.0 and 6.1, respectively), Pericytes (cluster 2) and Mesothelial cells (cluster 7), annotated based on differential gene expression analysis. In addition, several novel populations were identified that expressed modulators of WNT signaling, including genes encoding Secreted Frizzled Related Protein 1 (*SFRP1*) (SFRP1^hi^CEBPD+, cluster 1.0; SFRP1^hi^, cluster 1.1) and SFRP2+ (cluster 0). We also discovered a heterogenous population of cells enriched for expression of C-C Motif Chemokine Ligand 2 (*CCL2*): CCL2^hi^PRRX1+ (cluster 3.0), and CCL2^hi^CCL21+ (cluster 3.2). In contrast to the other mesenchymal populations, the mature VSM (0.35% of total mesenchymal cells at 8w; 2.9% at 10w) and CCL2^hi^CCL21+ (0.18% of total mesenchymal cells at 8w; 3% at 10w) populations expanded only after at 10w (**Figure S2A**), suggesting that their maturation occurs later in development compared to the rest of the mesenchymal compartment.

**Figure 2:**
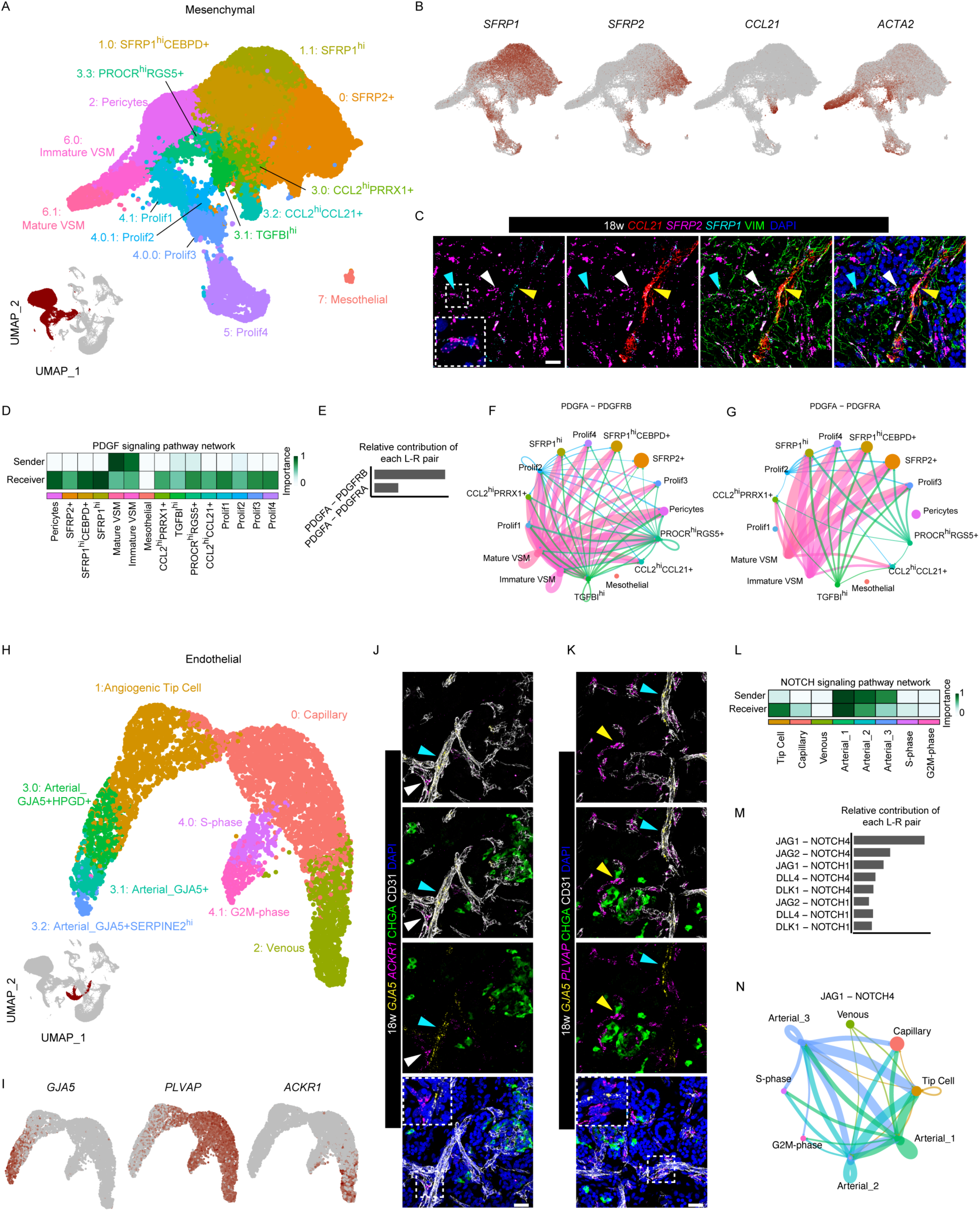
Identification of cell heterogeneity and cell-cell communication within the mesenchymal and endothelial lineages of the fetal human pancreas. (A) UMAP visualization of the cell populations comprising the human fetal pancreatic mesenchyme. (B) Feature plots depicting expression of *SFRP1, SFRP2, CCL21*, and *ACTA2* across the mesenchymal subpopulations. (C) 18 w human fetal tissue stained with an antibody against Vimentin (VIM; green) and ISH mRNA probes against *SFRP1* (cyan), *SFRP2* (magenta), *CCL21* (red), counterstained with DAPI (blue) to detect nuclei. Aqua, White, and Yellow arrowheads mark presumptive *SFRP1*+VIM+ SFRP1^hi^/SFRP1^hi^CEBPD+ cells, *SFRP2*+VIM+ SFRP2+ cells and *CCL21*+VIM+ CCL2^hi^CCL21+ cells, respectively. Dashed insets represent magnified regions. (D) Heatmap depicting predicted activity of the PDGF signaling pathway among the populations comprising the mesenchymal compartment. (E) Comparison of the predicted relative contribution of both significant PDGF signaling ligand receptor pairs, PDGFA-PDGFRB and PDGFA-PDGFRA, within the mesenchymal compartment as a whole. (F), (G) Circle plots depicting signaling mediated by PDGFA-PDGFRB (F) and PDGFA-PDGFRA (G) ligand-receptor pairs between cellular populations within the mesenchymal compartment. Line thickness is proportional to signaling strength, and line colors represent which population is the predicted “Sender” of the signal; colors match populations in (A). (H) UMAP visualization of the populations detected within the human fetal pancreatic endothelial Broad Group. (I) Feature plots depicting expression of *GJA5, PLVAP*, and *ACKR1* in the endothelial compartment. (J), (K) 18 w human fetal tissue stained with antibodies against pan-endothelial marker CD31 (gray) and pan-endocrine marker CHGA (green), along with ISH mRNA probes against (J) *GJA5* (yellow) and *ACKR1* (magenta) or (K) *GJA5* (yellow) and *PLVAP* (magenta); nuclei are counterstained with DAPI (blue). Aqua, white, and yellow arrowheads mark presumptive *GJA5*+CD31+ Arterial cells, *ACKR1*+CD31+ Venous cells and *PLVAP*+CD31+ Capillary/Venous cells, respectively. Dashed insets represent magnified regions. (L) Heatmap depicting predicted activity of the NOTCH signaling pathway among the various endothelial populations comprising the endothelial compartment. (M) Comparison of the predicted relative contribution of each significant NOTCH signaling ligand-receptor pair within the endothelial compartment as a whole. (N) Circle plot depicting the communication between endothelial populations with respect to JAG1-NOTCH4 signaling. Line thickness is proportional to signaling strength, and line colors represent which population is the “Sender” of the signal; colors match populations in (H). Scale bars are 25 um throughout.

We next validated the existence of novel mesenchymal subpopulations by performing multiplexed *in situ* hybridization (ISH) and immunofluorescence (IF) staining for *SFRP1*, *SFRP2,* and *CCL21* RNA and the broad mesenchymal protein marker VIMENTIN (VIM) at developmental stage 18 w **(Figure 2B-2C)**. *SFRP1*, *SFRP2*, and *CCL21* were found to colocalize with VIM, confirming their classification as mesenchymal. *SFRP1* was detected in a broad population of VIM+ mesenchymal cells, whereas *SFRP2* was expressed in a restricted subset of VIM+ cells that were also found to be positive for *SFRP1* (**Figure 2C**), confirming the predictions of the scRNA-Seq data (**Figure 2B**). In addition, *CCL21* expression overlapped with a subset of VIM+ mesenchyme (**Figure 2C**). Collectively, these data confirmed the *in vivo* existence of heterogeneous mesenchymal populations inferred by CellFindR.

To understand how mesenchymal cell subtypes may be communicating with one another via cell-cell signaling, we employed the computational package CellChat, which infers cellular communication within complex scRNA-Seq datasets (Jin et al., 2021). Signaling pathways scored as significantly active between mesenchymal subtypes included the FGF, EGF and COLLAGEN pathways. We focused on the PDGF signaling pathway, as it has been shown to be important for mesenchymal development in multiple other organs, including metanephric organs and the gastrointestinal tract (Wagner et al., 2007); (Karlsson et al., 2000). We found that the immature and mature VSM cell populations scored highest as producers of PDGF ligands (“Senders’’), while multiple clusters were denoted as “Receiver” populations (**Figure 2D**). In particular, the PDGFA-PDGFRB ligand-receptor pair was predicted to have the highest relative contribution to mesenchymal PDGF signaling, followed by PDGFA-PDGFRA (**Figure 2E**). To better understand the contribution of each ligand-receptor pair, we analyzed the dominant Senders and Receivers for each pair separately. For the PDGFA-PDGFRB ligand-receptor pair, we found that the dominant sources of PDGFA ligand were predicted to be the immature and mature VSM clusters, and the dominant receiver through the PDGFRB receptor was predicted to be pericytes (**Figure 2F**). This result is similar to PDGF signaling in the retina, where pericytes are recruited to developing endothelium through PDGF signaling to aid in formation of the blood-retinal barrier (Park et al., 2017). When analyzing the PDGFA-PDGFRA ligand receptor pair, we found that the dominant receivers of the PDGFA ligand through the PDGFRA receptor were predicted to be the SFRP-expressing clusters (SFRP1^hi^CEBPD+, SFRP1^hi^, and SFRP2+) (**Figure 2G**). Currently, the effects of PDGFA-PDGFRA signaling in these populations are unknown and warrant further studies.

Endothelial cell-derived signals are essential for proper formation of the murine pancreas (Lammert et al., 2001, 2003). The murine pancreas is surrounded by vasculature by as early as E10.5, with arterial and venous specification occurring at E11 (Azizoglu et al., 2016). In human fetal pancreas tissue, CD31+ blood vessels are present as early as 7 w (Roost et al., 2014). Previous studies demonstrated phenotypic (Henderson and Moss, 1985) and functional (Azizoglu et al., 2016; Zanone et al., 2008) heterogeneity within endothelial populations. However, transcriptional heterogeneity among endothelial cells in the human fetal pancreas has not yet been investigated at the single cell level. Thus, we next focused on the endothelial compartment of our single-cell dataset to better understand its developmental role in pancreatic organogenesis. CellFindR identified eight clusters within the endothelial Broad Group, with five main subtypes (**Figure 1E**). This included *RGCC*+ capillaries (cluster 0), *NR2F2+/ACKR1+* venous cells (cluster 2), *PRND+/IGFBP3+* angiogenic tip cells (cluster 1), proliferating cells (clusters 4.0 and 4.1), and a heterogeneous population of *GJA5+* arterial cells (*GJA5+/HPGD+*; cluster 3.0; *GJA5+*; cluster 3.1; *GJA5+/SERPINE2^h^*^i^, cluster 3.2) (**Figure 2H**). We detected no major shifts in prevalence of these populations across the developmental time points within our dataset, suggesting that pancreatic endothelial specialization is already established at 8 w (**data not shown**).

To validate our CellFindR inferences, we performed ISH for arterial marker *GJA5* (Buschmann et al., 2010), capillary/venous marker *PLVAP* (Guo et al., 2016) and venous marker *ACKR1,* along with antibody staining for pan-endothelial marker CD31 in 18 w fetal tissue (**Figure 2I-2K**). *In vivo* analysis revealed mutually exclusive *GJA5*+ arterial and *ACKR1*+ venous blood vessels that co-localized with CD31+ (**Figure 2I**). We also observed *PLVAP*+/CD31+ capillary/venous cells that did not colocalize with *GJA5* probe (**Figure 2K**). Taken together, these data explore and confirm endothelial cellular heterogeneity in the developing fetal pancreas.

When investigating cellular communication among pancreatic endothelial populations, we found that NOTCH signaling scored among the highest pathways in our CellChat analysis. NOTCH signaling has previously been shown to be a critical regulator of endothelial specification (Akil et al., 2021) but its role in fetal pancreatic endothelium has yet to be described. We found that the arterial populations 1, 2, and 3 scored the highest as “Senders” of NOTCH ligands, while the Arterial_1 and Angiogenic Tip Cell populations scored the highest as “Receivers” (**Figure 2L**). When assessing the contribution of each ligand-receptor pair, we observed that the JAGGED1(JAG1)-NOTCH4 pair was predicted to make the highest contribution to NOTCH signaling within our endothelial dataset (**Figure 2M**). NOTCH ligand JAG1 has been implicated as a pro-angiogenic molecule that is capable of counteracting the anti-angiogenic effects of Delta-Like Canonical Notch Ligand 4 (DLL4)-NOTCH signaling in mice (Benedito et al., 2009). Within our dataset, the arterial populations were scored highest as “Senders” of the JAG1 ligand, with both the Angiogenic Tip cell and Arterial_1 population predicted as “Receivers” (**Figure 2N**). These data suggest that pancreatic arterial cells may maintain angiogenic tip cell fate through NOTCH signaling mediated by the pro-angiogenic molecule JAG1.

Heterogeneity was also discovered within the remaining Broad Groups. In the exocrine compartment, for instance, we observed *CFTR*-expressing ductal cells (clusters 2.0, 2.1.0 and 2.1.1) (Hyde et al., 1997). In addition, three sub-clusters of acinar cells appeared to represent a continuum of maturation states, characterized by varying degrees of expression of digestive enzymes (clusters 1, 3.0, and 3.1) (**Figure S2B**). We annotated Exocrine clusters 0.0 and 0.1 as pre-acinar and pre-ductal cells, respectively, based on their displaying gene expression profiles most highly correlated to acinar and ductal cells, respectively (**data not shown**); these may represent tip and trunk cells that eventually give rise to the differentiated exocrine tissue (Zhou et al., 2007). In the Neuronal Broad Group, we identified clusters representing myelinating Schwann Cells (clusters 0.0, 0.1, and 0.2), as well as peripheral nerve subtypes expressing various neurotransmitters such as *VIP, NPY* and *NOS1* (clusters 1.0.0, 1.0.1, 1.1.0, 1.1.1 and 1.2) and proliferating neuronal cells (clusters 2.0.0, 2.0.1, 2.1.0, 2.1.1, 2.2) (**Figure S2C**). In the Immune Broad Group, we identified 16 lymphoid lineage populations, including T cells, B cells, and NK cells. We also identified 15 myeloid lineage populations, including neutrophils, monocytes, macrophages, and mast cells (**Figure S2D**). The proportion of myeloid and lymphoid lineage cells remained relatively stable from 8 w to 12 w, while a substantial increase in the proportion of lymphoid cells was observed at 16 w, possibly due to the infiltration of blood B cells **(data not shown)**.

Next, we deployed the CellChat algorithm (Jin et al., 2021) to interrogate the incoming and outgoing signaling pathways active among all subtypes of each Broad Group **(Figure S2E-S2F)**. The Midkine (MK), pleiotrophin (PTN), and RESISTIN pathways were the top 3 most highly scored both for the incoming and the outgoing signaling. Previous studies in mice revealed the fundamental function of VEGF signaling in islet vascularization and vessel architecture (Azizoglu et al., 2016; Zanone et al., 2008) CellChat predicted that in the developing human fetal pancreata, VEGF signaling was strictly sensed by endothelial cells (receivers), and secreted by epithelial cells and VEGF signaling cross talk was also observed between pancreatic mesenchyme, epithelium (senders) and endothelial cells (receivers), suggesting that mesenchymal and epithelial cells promote vascular modeling through the secretion of VEGF in the fetal human pancreas, paralleling murine development. In addition, mesenchymal cells were found to produce a variety of paracrine factors, including WNT and FGF ligands **(Figure S2F)**, which are critical for endothelial or epithelial development (Ye et al., 2005). CXCL signaling secreted from the endothelial Broad Group was predicted to act on a wide range of immune cell types, including monocytes, macrophages, and DCs **(Figure S2E-S2F)**. Taken together, these coordinated signaling interactions across cell types attest to the significance of cell-to-cell communication in human fetal pancreas organogenesis and provide a framework for future studies involving heterotypic cellular signaling in the human fetal pancreas.

### Discovery and Characterization of Novel Human Endocrine Progenitor Cell Populations

Diabetes mellitus is one of the most common endocrine disorders worldwide, affecting hundreds of millions of individuals across the globe (Saeedi et al., 2019). Mapping the cellular and molecular landscape during human endocrine development is a critical step for improving stem cell-derived therapies for diabetes. CellFindR permitted the identification of four hormone-expressing endocrine clusters in the human fetal pancreas, distinguished by the expression of *INS* in beta cells (cluster 0; 6,700 cells), *GCG* in alpha cells (cluster 1; 3,388 cells), *SST* in delta cells (cluster 3; 1,554 cells), and *GHRL* in epsilon cells (cluster 4; 909 cells) (**Figure 3A-B**).

**Figure 3:**
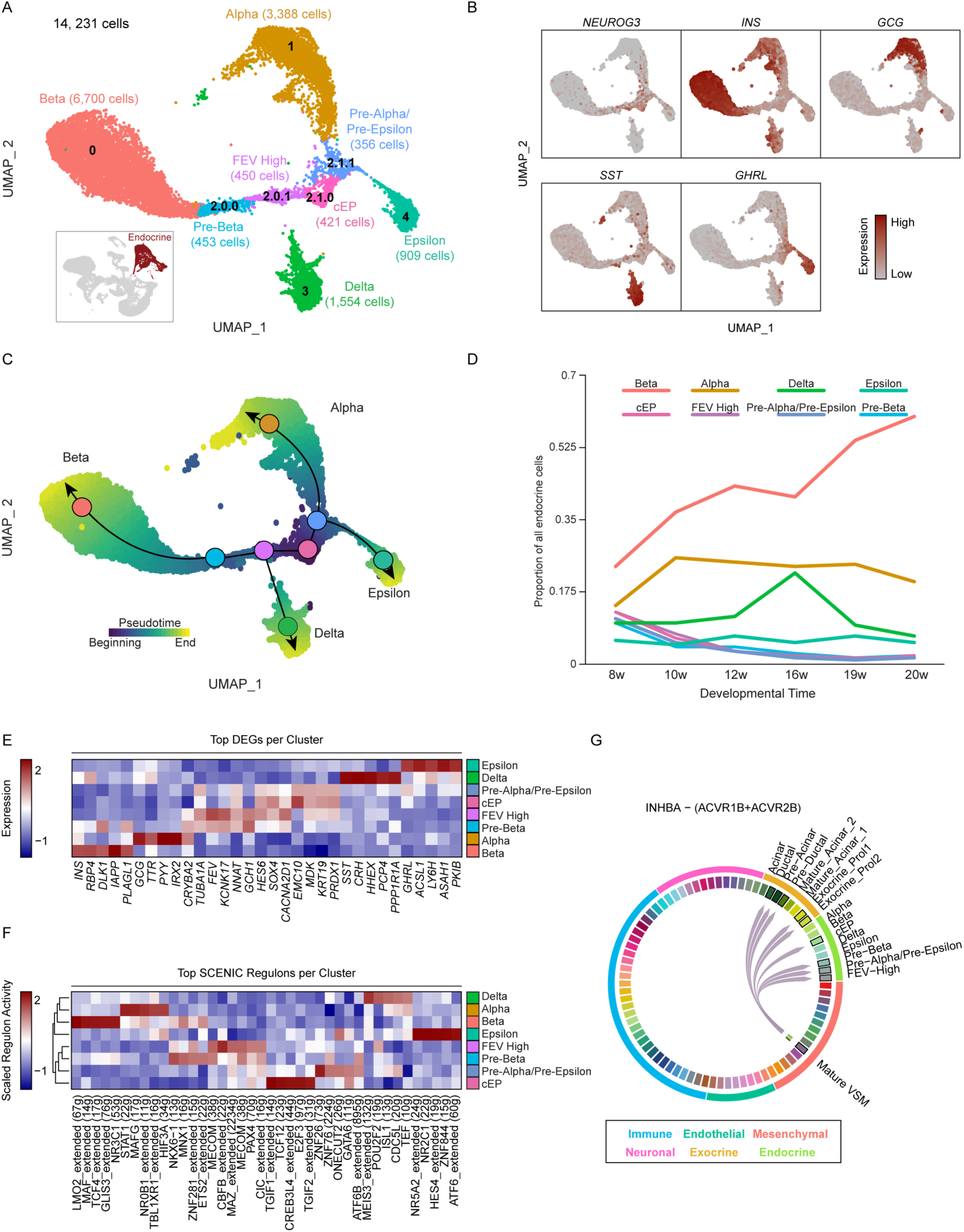
Discovery of four novel putative progenitor populations and unique lineage dynamics in the developing human endocrine pancreas. (A) UMAP visualization of sub-clustered endocrine populations identified in the merged dataset as shown in Figure 1B (inset). (B) Feature plots show expression of known markers of endocrine cell types, including *NEUROG3* to mark endocrine progenitors, *INS* to mark beta cells, *GCG* to mark alpha cells, *SST* to mark delta cells, and *GHRL* to mark epsilon cells. (C) Pseudotime reconstruction of endocrine lineage trajectories assembled using Slingshot, with the centroid of each cluster along the lineage depicted with a circle. (D) Line graph showing the representation of each endocrine cell population as a proportion of the total number of endocrine cells, across developmental time. (E) Heatmap depicting expression levels of the top 5 differentially-expressed genes per endocrine cluster. (F) Heatmap depicting the scaled regulon activity of the top SCENIC regulons per cluster. (G) Chord diagram depicting predicted INHBA-ACVR1B/ACVR2B signaling among the human fetal pancreas dataset. Cell populations in the heatmaps in (E) and (F) are colored to match clusters in (A).

CellFindR also annotated a remaining fifth endocrine cluster, cluster 2, which we classified as a putative endocrine progenitor (EP) cell population based on its specific expression of *NEUROG3* and lack of hormone expression (**Figure 3A-3B**). CellFindR further sub-clustered cluster 2 into four sub-clusters: 2.0.0 (453 cells), 2.0.1 (450 cells), 2.1.0 (421 cells), and 2.1.1 (356 cells), resulting in a final endocrine dataset comprised of eight clusters (**Figure 1H**, **Figure 3A**). These results reveal heterogeneity within the progenitor pool of the human pancreatic endocrine compartment.

To test the hypothesis that the four subclusters of cluster 2 indeed represented biologically distinct endocrine progenitor states, we utilized the R package Slingshot (Street et al., 2018) to perform lineage reconstruction and pseudotemporal ordering of all endocrine cells from all developmental timepoints (**Figure 3C**). Lineage reconstruction revealed a bifurcated trajectory in which cells from cluster 2.1.0 were ordered at the beginning of pseudotime; we henceforth refer to this cluster as the Common Endocrine Progenitor (cEP) population. Cluster 2.1.1 (henceforth referred to as the Pre-Alpha/Pre-Epsilon population) followed the cEP cluster in pseudotime and itself served as a further bifurcation point, leading to either the Alpha Cell or Epsilon Cell clusters (**Figure 3A**, **3C**). Along an alternative branch, cEP cells were followed in pseudotime by cluster 2.0.1, which expressed the highest levels of the gene *FEV* (**Figure 3C, 3E**), a transcription factor we and others previously identified as a marker of a novel EP cell state during murine endocrine development (Bastidas-Ponce et al., 2019; Byrnes et al., 2018; Yu et al., 2019b). This highly *FEV*-expressing cluster, henceforth referred to as the FEV High population, was followed in pseudotime by cluster 2.0.0 (henceforth referred to as the Pre-Beta population), terminating at the Beta Cell cluster. The reconstruction by Slingshot positioned the FEV High population as a progenitor of delta cells, as well as beta cells (**Figure 3C**). Additionally, Pearson correlation analysis among the endocrine populations revealed high correlation of the EP populations cEP, Pre-Alpha/Pre-Epsilon, and FEV High with one another, while the Pre-Beta population was closest correlated with the Beta population (**Figure S3B**). These data predict that the lineage trajectories in human endocrine development are distinct from those reported in mouse and human development (Bastidas-Ponce et al., 2019; Byrnes et al., 2018; Yu et al., 2019b, 2021)

To assess how each of these distinct endocrine cell states varied across developmental time, we quantified their population dynamics across developmental time. Each of the four EP populations was present in every biological scRNA-seq sample, across all timepoints sampled (**Figure S3A**). The relative proportion of each of the EP populations was highest at 8 w (cEP, 12.8%; Pre-Alpha/Pre-Epsilon, 11.0%; FEV High, 12.4%; Pre-Beta, 10.2% of all endocrine cells), then decreased as developmental time progressed to reach their lowest levels at 20 w (cEP, 2.2%; Pre-Alpha/Pre-Epsilon, 1.6%; FEV High, 1.8%; Pre-Beta, 2.1% of total endocrine cells) (**Figure 3D**), consistent with the model that they represent progenitor populations. In contrast, the proportion of beta cells in the developing pancreas steadily increased across developmental time (24% of all endocrine cells at 8 w; 60% at 20 w). The proportion of alpha, delta and epsilon cells remained relatively stable from 10 to 20 w at proportions approximating those reported in human adult islets (Da Silva Xavier, 2018). These data are consistent with a model whereby the pool of more differentiated endocrine cells increases over developmental time at the expense of a dwindling progenitor pool. These data further support the hypothesis that the four endocrine cell subclusters represent novel endocrine progenitor populations in the human fetal pancreas.

### Transcriptional Regulation of Human Pancreatic Endocrine Development

To identify the transcriptional features that distinguish each endocrine cell population, we performed a series of analyses. First, we conducted differential gene expression analysis across all eight endocrine clusters by comparing each cluster against all other clusters, resulting in a total of 858 genes with at least 0.5 log2 fold change in expression (**Figure S3C**). Of these 858 differentially expressed genes (DEGs), the majority were most highly expressed by the EP populations (643 genes). Pre-Beta (n = 73 genes) and FEV High (n = 163 genes) clusters expressed genes associated with neuronal development (*TUB1A1, NNAT* by Pre-Beta; *FEV* by FEV High) (Aiken et al., 2017; Kanno et al., 2019; Krueger and Deneris, 2008), while the cEP cluster expressed genes (n = 239) associated with mRNA processing and chromatin remodeling (*SRSF3, RBMX*) (Ajiro et al., 2016; Zhou et al., 2019), potentially representing dynamic priming of gene expression needed for endocrine differentiation. Pre-Alpha/Pre-Epsilon cells also expressed genes (n = 168) associated with neuronal development and RNA processing (**Figure S3C**). Pathway analysis of the differentially-expressed genes corroborated these findings, revealing that enriched pathways included those annotated as being involved in neuronal development (Pre-Alpha/Pre-Epsilon) and endocrine function (Pre-Beta) (**Figure S3D**).

Among hormone-producing cells, DEGs expressed by the Beta Cell population (n = 27 genes) included genes poorly characterized in beta cells (*LMO2*, *ASPH*), as well as known markers of beta cell identity (*INS)* (Wang et al., 2007) and function (*IAPP, SLC30A8, PCSK1*) (Pound et al., 2009); (Taylor et al., 2020); (Ramzy et al., 2020) (**Figure 3E**). Alpha Cells (n = 37 genes) expressed both known alpha cell-related genes, (*IRX2*, *TTR*), as well as genes whose function in alpha cells is not well characterized, such as *EDN3*, *SPINT2,* and *CDNK1C*. Delta Cells (n = 23 genes) highly expressed markers of delta cell identity (*SST*) and genes encoding peptides with known endocrine function (*NPW, CRH*) (Mondal et al., 2006); (Childs et al., 1995) (**Figure 3E**). Of the four hormone-expressing cell types, Epsilon Cells had the highest number of DEGs detected (n = 127 genes), including genes associated with epsilon cell identity and function (*GHRL*), and genes involved in cellular signaling (*FGF12, FGF1)* (**Figure 3E**). The enriched pathways in the hormone-expressing populations included Insulin Processing (Beta Cells), Retinoic Acid Signaling (Alpha Cells) and ATF-2 Transcription Factor Network signaling (Epsilon Cells) (**Figure S3D**). Together, these data describe differentially expressed transcriptional programs and enriched signaling pathways within each human pancreatic endocrine population. We next set out to focus specifically on differentially-expressed transcription factors (TFs), as they are critical regulators of cell fate determination (Conrad et al., 2014; Jennings et al., 2015). We identified 108 TFs that were differentially expressed across the endocrine lineage (**Figure S3C**). cEP cells displayed highest expression of *NEUROG3* targets *NKX2-2*, *NEUROD1*, and *INSM1* (Breslin et al., 2007; Churchill et al., 2017; Gasa et al., 2008), as well as TFs involved in Hippo-YAP signaling (*TEAD2*) and Notch signaling (*RBPJ* and *HES6*), consistent with previous evidence in mice that Notch activity is critical for maintenance of endocrine progenitor cell fate (Murtaugh et al., 2003) (Figure S3C). The cross-inhibitory interactions between *Pax4* and *Arx* promote the acquisition of beta cell or alpha cell fate during murine endocrine development (Collombat et al., 2003). Similarly, we found that the FEV High and Pre-Beta EP populations expressed high levels of *PAX4*, whereas the Pre-Alpha/Pre-Epsilon EP cluster showed elevated levels of *ARX* expression during human pancreas development. Expression of the TF *NKX6.1*, a crucial regulator of beta cell fate **(Schaffer et al., 2013)**, was enriched in the FEV High and Pre-Beta populations. Moreover, the highest expression of beta cell regulators *PDX1 (Gao et al., 2014)*, *MNX1 (Pan et al., 2015)*, and *PAX6 (Swisa et al., 2017)* appeared in the Pre-Beta but not in the FEV High EP cells, suggesting that these genes may play a functional role in beta cell fate restriction. We also identified TFs that were specifically expressed in a specific EP cell type, such as *TOX3* (cEP)*, SIM1* (Pre-Beta), and *POU2F2* (Pre-Alpha/Pre-Epsilon) **(Figure S3C)**. These results provide a rich dataset of transcription factors that warrant further study in developing human endocrine cells.

We next aimed to compare the gene expression profiles among the four sub-clustered EP populations to identify which genes had driven their distinction by CellFindR. Pairwise comparison of Pre-Beta vs. FEV High cells revealed higher expression of genes associated with beta cell maturation and function (*MAFB, PCSK1N, GNAS*) in the Pre-Beta cluster, while the FEV High cluster showed higher expression of genes such as *FEV* and *HES6* (**Figure S3E**). Comparison of the cEP vs. Pre-Alpha/Pre-Epsilon clusters uncovered higher expression in cEP cells of genes such as *SOX4,* which cooperates with *Neurog3* to regulate endocrine induction in the murine pancreas (Xu et al., 2015). In contrast, Pre-Alpha/Pre-Epsilon cells more highly expressed genes associated with differentiated endocrine cells, such as *ARX*, *ISL1* and *GHRL*. The results from these pairwise comparisons corroborate our hypothesis, based on lineage reconstruction, that the EP populations represent distinct cell states that are pre-committed to one or more hormone-producing cell fates. Taken together, these data have enabled the construction of a model of human endocrine lineage specification and the identification of novel genes governing cell fate decisions during human fetal development.

### Elucidating Active Regulons Governing Endocrine Cell Fate

To identify TFs that show functional evidence of characteristic downstream activity within distinct cell populations, we utilized the R package SCENIC (Single Cell rEgulatory Network Inference and Clustering). This method scores TF activity based on the collective expression of a given TF and its direct gene targets (together referred to as a “regulon”), thus identifying at single-cell resolution the gene regulatory networks (GRNs) that are likely to govern human endocrine cell fate decisions (Aibar et al., 2017). SCENIC analysis identified a total of 256 active regulons across the eight endocrine populations, with hierarchical clustering grouping the hormone-expressing populations as most similar to one another and the EP populations most similar to one another based on regulon activity **(Figure S3F)**. Specifically, highly scored regulons consisted of known regulators governing endocrine differentiation, such as the MAF family of TFs (*MAF*, Beta cells; *MAFG*, Alpha cells), *NKX6.1* (Pre-Beta), and *ISL1* (Delta) (**Figure 3F**). Importantly, SCENIC also determined active regulons not present in our original TF list generated by DE analysis. These active regulons included those involved in WNT signaling (*TCF4*, Beta cells; *TCF12*, cEP) as well as the ETS family of TFs (*ETS1,* Epsilon; *ETS2,* Pre-Beta). We found that the activity score of the *FEV* regulon was the highest in the FEV High population, pointing to a potential functional role of the TF *FEV* in regulating beta cell fate. Taken together, our data predict “active” TFs (regulons) governing human pancreatic endocrine development.

### Cell-Cell Communication Among Human Endocrine Cells

Heterotypic cellular signaling, mediated by factors such as WNT, FGF and EGF, is essential for the proper formation of epithelial cells in the murine pancreas (Bhushan et al., 2001; Miettinen et al., 2000; Sharon et al., 2019). We further analyzed the signaling pathways identified by CellChat to determine which might play a role in human endocrinogenesis. We observed little evidence of cellular signaling incoming to the endocrine compartment through the WNT, FGF, or EGF signaling pathways (**Figure S2D**). However, ACTIVIN signaling was specifically activated in the epithelium, including the four EP populations in the Endocrine Broad Group, as well as the Pre-Acinar, Pre-Ductal and proliferating exocrine populations in the Exocrine Broad Group **(Figure S2D).** Given the role of Activin signaling in guiding pancreatic morphogenesis and endocrine differentiation in mice (Zhang et al., 2004), we next investigated which ACTIVIN ligand-receptor pair contributed to signaling within the fetal human pancreas. We found that interaction between INHBA (an ACTIVIN ligand) and ACVR1B/ACVR2B (ACTIVIN receptors) was the sole ligand-receptor pair identified as significant (**Figure 3G**). Unexpectedly, among all 103 cell types in the developing human pancreata, the Mature VSM cluster within the mesenchymal compartment was identified as serving as the sole source of INHBA ligand, suggesting that Mature VSM plays a role in human fetal endocrine differentiation (**Figure S2E, 3G**). Our results predict that pancreatic endocrinogenesis in humans depends on signal input from the mesenchymal niche environment.

### Confirmation of Novel Endocrine Progenitor Populations *In Vivo*

To confirm the findings of EP cell heterogeneity that had emerged from the scRNA-Seq data, we first set out to identify genes or combinations of genes that could serve as specific markers of each of the four EP populations. Manual curation of top differentially-expressed genes across EP cells identified Sushi Domain Containing 2 (*SUSD2),* LIM Homeobox Transcription Factor 1 Beta *(LMX1B)*, Peripherin (*PRPH*), and Aristaless-Related Homeobox *(ARX)* as highly enriched in the cEP, FEV High, Pre-Beta, and Pre-Alpha/Pre-Epsilon EP populations, respectively **(Figure 4A-4B).** Among these four genes, *SUSD2* has previously been broadly described to label *NEUROG3* expressing EP cells in the developing human pancreas (Liu et al., 2014; Ramond et al., 2017)*. ARX* is an important regulator of alpha cell fate (Itoh et al., 2010) that has not been described to mark human Pre-Alpha/Pre-Epsilon EP cells. *LMX1B* and *PRPH* are novel markers labeling sequential progenitor states during human beta cell development.

**Figure 4:**
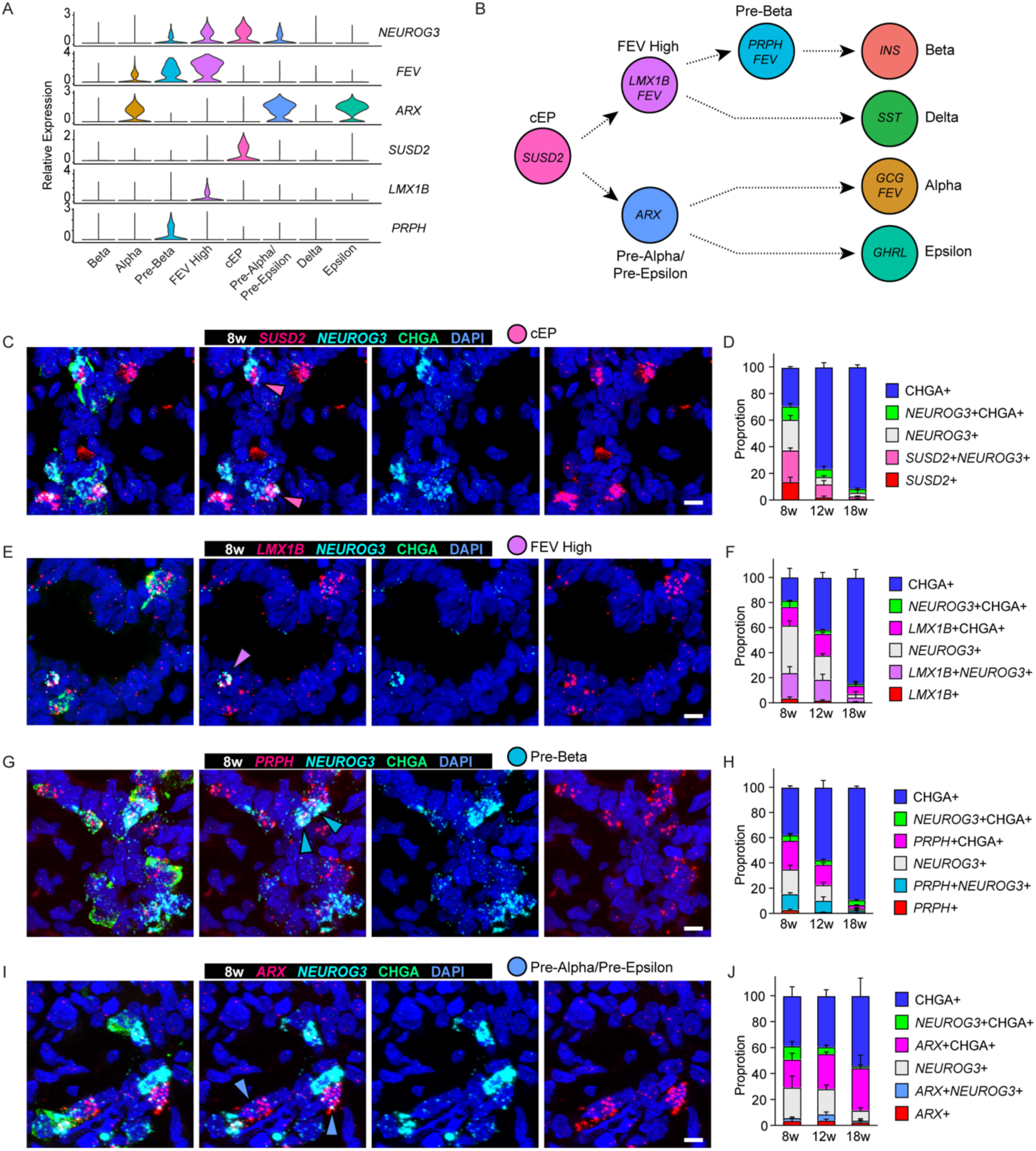
*In vivo* confirmation of novel endocrine progenitor cell populations. (A) Violin plot depicting expression of endocrine progenitor (EP) marker genes. (B) Model showing predicted lineage relationships among developing human endocrine cells, along with genes that mark each population. (C) (E) (G) (I) 8 w human fetal tissues were stained with an antibody against the differentiated endocrine cell marker CHGA (green), along with probes against *NEUROG3* (cyan), and putative EP cell markers (red) (C) *SUSD2*; (E) *LMX1B*; (G) *PRPH*; (I) *ARX* for detection by *in situ* hybridization (ISH). Nuclei were counterstained with DAPI (blue). Scale bars, 10 um. Pink, Purple, Aqua, and Blue arrowheads mark presumptive *SUSD2*+*NEUROG3*+CHGA-cEP, presumptive *LMX1B*+*NEUROG3*+CHGA+ FEV High EPs, presumptive *PRPH*+*NEUROG3*+CHGA+ Pre-Beta EPs, and presumptive *ARX*+*NEUROG3*+CHGA+ Pre-Alpha/Pre-Epsilon EPs, respectively. (D) (F) (H) (J) Quantification of staining as performed in (C), (E), (G), (I) extended across a time course of 8, 12, and 18 w. Y-axis represents the proportion of cells positive for at least one of the makers. Graphs are presented as mean ± SEM (n=3 biological replicates).

Once markers of each of the four novel EP subtypes had been identified, we next validated the existence of each EP cell state *in vivo* by performing multiplexed *in situ* hybridization (ISH) and immunofluorescence (IF) staining on independent human fetal pancreata at 8, 12, and 18 w. ISH was performed for each putative marker gene alongside the pan-EP marker *NEUROG3* and combined with IF staining for the pan-differentiated endocrine cell marker CHGA. When staining tissues to validate the presence of the cEP population, we found five unique cell states based on their expression with the combination *SUSD2*/*NEUROG3*/CHGA (**Figure 4C-4D)**. As predicted by our scRNA-Seq analysis, we detected putative cEP cells, characterized as *SUSD2^+^NEUROG3*^+^CHGA^-^ (23.7% at 8w, 9.8% at 12w, 2.1% at 18w) **(Figure 4C-4D)**. The same experimental and quantification approaches were adopted to validate the existence of the other three EP types. Thus, we observed the presence of *LMX1B^+^NEUROG3*^+^ (putative FEV High EP cells; 20.2% at 8 w, 16.7% at 12 w, 3.6% at 18 w), *PRPH*^+^*NEUROG3^+^* (putative Pre-Beta EP cells; 12.6% at 8 w, 9.3% at 12 w, 2.5% at 18 w), and *ARX*^+^*NEUROG3*^+^ (putative Pre-Alpha/Pre-Epsilon EP cells; 2.2% at 8 w, 4.9% at 12 w, 1.1% at 18 w) cells that were also negative for CHGA **(Figure 4E-4J)**. All four EP populations were detected in nine independent biological samples of pancreas tissue.

To assess the exclusivity of the four putative endocrine progenitor populations, we next performed multiplexed ISH staining at 8w to detect markers of all four EP cell types simultaneously; we found the expression of genes predicted to specifically mark each EP population indeed showed mutual exclusivity *in situ* **(Figure S4A)**. Moreover, the relative prevalence of all four EP populations decreased over developmental time **(Figure 4D, F, H, J)**, consistent with the characterization of these populations as progenitor populations. Collectively, these results confirmed the presence of the novel EP subtypes in the developing human pancreas as predicted by computational analysis of the scRNA-seq data.

Our CellFindR inferences identified *FEV* as a marker for both FEV High and Pre-Beta EP populations (**Figure 4A**). Given that we previously had identified *Fev* as a marker of endocrine progenitor cells in the murine pancreas (Byrnes et al., 2018; Yu et al., 2019b), we assessed the dynamics of *FEV* expression in the human fetal pancreas. Consistent with our scRNA-Seq data, the percentage of *FEV^+^NEUROG3^+^* double positive cells (marking FEV High or Pre-Beta EP cells) decreased over developmental time (13.4% at 8 w, 5.9% at 12 w, 2.8% at 18 w) (**Figure S4B**). We also observed a significant rise in *FEV^+^*CHGA^+^ double positive cells in 18 w tissue compared to 12 w and in 12 w tissue compared to 8w **(Figure S4B)**. Further investigation revealed that *FEV* expression localized to GCG-expressing alpha cells, but not INS*-*expressing beta cells, at 18 w **(Figure S4C).** This observation agrees with previously published reports that in adult human islets, *FEV* is exclusively expressed in the alpha cells (Camunas-Soler et al., 2020; Muraro et al., 2016; Segerstolpe et al., 2016). Along the beta cell lineage trajectory, *FEV* expression was detected in the FEV High and Pre-Beta populations, but absent in differentiated beta cells themselves. In contrast, along the alpha cell lineage, *FEV* was expressed in differentiated alpha cells themselves but not in their progenitors **(Figure 4A-4B, S4D)**. These results demonstrate regulation of *FEV* expression during human endocrine development and suggest a dynamic and lineage-specific role in regulating alpha vs. beta cell fate and/or function.

### Single-nucleus ATAC-Seq of Human Fetal Endocrine Cells Reveals Dynamic Chromatin Accessibility

Recent advances in single-cell technologies have allowed for the integration of multi-omic single-cell data, leading to new insights into developmental biology (Buenrostro et al., 2018; Lake et al., 2018; Ranzoni et al., 2021). We set out to gain an understanding of the epigenetic mechanisms upstream of gene expression that are important in governing cell identity during endocrine development. To this end, we performed snATAC-Seq on human fetal pancreas using the 10x Genomics Chromium Next GEM Single Cell ATAC v1.1 platform. To increase the resolution for endocrine cell types, we enriched for EPCAM+ epithelial cells in 12-week fetal pancreas, a particularly active time of cell expansion and diversification (**Figure 5A**). Filtering, dimensional reduction, initial clustering, and analysis steps were performed with the R package ArchR(Granja et al., 2020), resulting in a final dataset comprising 6,010 nuclei with a median of 31,835 fragments captured per nucleus **(data not shown).** Among these nuclei, 1,754 were classified as belonging to endocrine cells (**Figure 5A**) based on the Gene Score Matrix (accessibility of gene body plus promoter) of *CHGA*.

**Figure 5.**
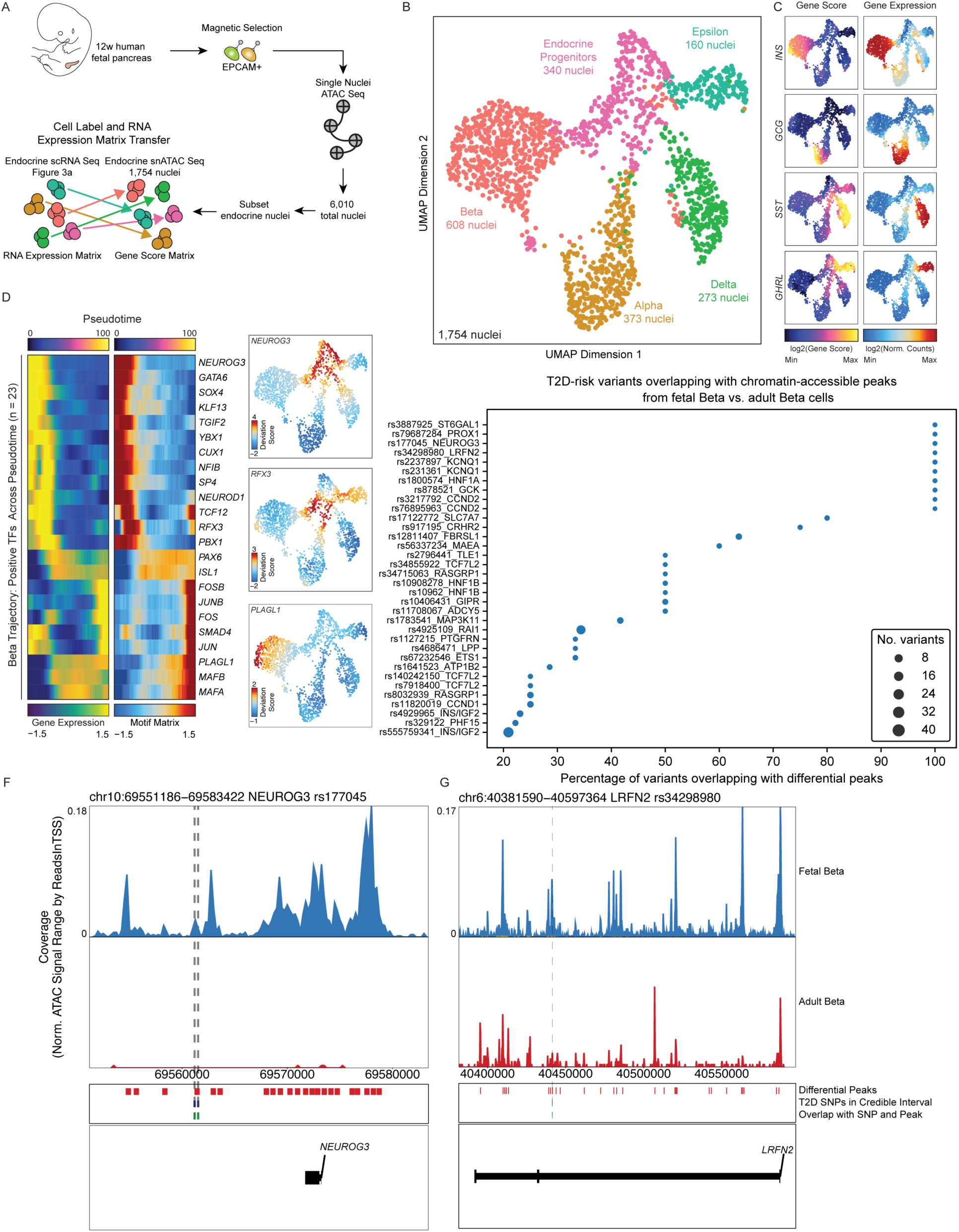
Single-nucleus ATAC-Seq reveals chromatin accessibility dynamics and type 2 diabetes genetic risk loci in the developing human endocrine pancreas. (A) Schematic of workflow for single-nucleus ATAC-Seq (snATAC-Seq) performed on EPCAM+ enriched cells from 12w human fetal pancreas. (B) UMAP of endocrine snATAC-Seq data reveals populations of hormone-expressing cells (alpha, beta, delta, and epsilon cells) as well as endocrine progenitors (collapsed here into a single population; for endocrine progenitors broken down into 4 subpopulations see Extended Figure 5A). (C) Feature plots showing ATAC gene scores (left) and corresponding RNA expression values (right) from integration of snATAC-Seq/scRNA-Seq data. (D) Heatmaps depicting gene expression levels (left heatmap) and motif enrichment scores (right heatmap) of positive transcription factors (those with correlated gene expression and motif enrichment) along the beta cell lineage. Motif deviation scores for selected transcription factors are displayed at single-cell resolution in the feature plots to the right. (E) T2D-risk loci enriched in differentially accessible peaks of fetal beta cells vs. adult beta cells. Scatter size correlates with the number of SNPs in the genetic credible interval. (F), (G) Track plots displaying accessibility of the *NEUROG3* and *LRFN2* loci in fetal (top) vs. adult (bottom) beta cells. Differential peaks, T2D risk variants in the genetic credible interval, and the T2D risk variants overlapping with differential peaks were highlighted.

Next, we integrated our endocrine snATAC-Seq and scRNA-Seq datasets to perform multi-omic analysis in the same cell types. Each cell in the snATAC-Seq dataset was correlated with its most similar counterpart in the scRNA-Seq dataset by correlating the Gene Score Matrix with the scRNA-Seq expression matrix (RNA transcript counts) on a per-cell basis (**Figure 5A**). Once these highly correlated pairs were found, the snATAC-Seq data from each cell were associated with the corresponding cell type label and RNA expression matrix. Of note, transfer of a cell label was not forced if the inferred gene score from snATAC-Seq data did not correlate with gene expression in any of the cells within the scRNA-Seq dataset. Our integration analyses allowed the identification of eight endocrine populations in the snATAC-Seq dataset, including the four newly identified EP populations (cEP, FEV High, Pre-Beta, and Pre-Alpha/Pre-Epsilon) and four hormone-expressing populations (**Figure S5A**). The Gene Score of the EP marker genes identified by scRNA-Seq analysis showed high concordance with their corresponding RNA expression levels (**Figure S5B**), confirming the existence of EP cell states using single nucleus chromatin accessibility analysis. Due to the low numbers of nuclei within each individual EP subpopulation, we merged all four of the EP populations into one single cluster, resulting in a final dataset of 5 clusters, consisting of Alpha (373 nuclei), Beta (608 nuclei), Delta (273 nuclei), Epsilon (160 nuclei), and pooled Endocrine Progenitor cells (340 nuclei) (**Figure 5B**). These data provide further confirmation of the discovery by scRNA-seq of four EP cell states, using an orthogonal method of snATAC-Seq.

To identify regions of accessible chromatin in the various cell types, we used the peak calling algorithm MACS2(Zhang et al., 2008). Among the 190,995 peaks aggregated across all five populations, 40,635 peaks were differentially accessible on a per-cluster basis (**data not shown**). As expected, the local chromatin of hormone genes *INS, GCG, SST,* and *GHRL* exhibited differential accessibility across all populations (**Figure 5C**). To calculate motif deviations (predicted TF activity) on a per-cell basis across all endocrine cells, we used chromVAR(Schep et al., 2017) and then correlated the deviation scores with the integrated RNA expression matrix to identify “positive” TFs for which expression was highly correlated with motif accessibility. This analysis identified a list of 49 “positive” TFs (**Figure S5C**), including known regulators of endocrine differentiation such as *NEUROG3, MAFA,* TCF family TFs (*TCF3, TCF4, TCF12, TCF15*) and *PDX1*, as well as TFs that warrant further functional studies, such as *PLAGL1* and *CUX1*. We then compared the 49 “positive” TFs from this multi-omic analysis with the 253 “active” TFs previously inferred by SCENIC analysis. We found that 21 of the TFs were overlapping, including *MAFA*, *TCF4*, and *RFX5,* suggesting critical regulatory roles in the developing human pancreas **(data not shown)**.

To identify transcription factors across the endocrine lineages that might be governing cell fate decisions, we performed lineage trajectory inference among the snATAC-Seq endocrine populations and applied the same “positive” TFs determination method as above along each lineage (**Figure 5D**). We observed robust changes in chromatin accessibility across the Beta, Alpha, Delta, and Epsilon trajectories, and identified “positive” TFs that may drive the specification of each hormone-expressing endocrine cell population (e.g., PLAGL1 in the Beta lineage). Taken together, our multi-omic analyses identified TFs with potential functions in orchestrating chromatin accessibility and mediating endocrine differentiation.

### Identification of Development-specific Type 2 Diabetes GWAS Risk Loci

In contrast to Type 1 Diabetes (T1D) were the majority of genetic association signals exert their effect through the immune system (Kim et al., 2021), there is compelling evidence fromphysiology (De Franco, 2020; Dimas et al., 2014) and epigenomics (Thurner et al., 2018) that pancreatic islets are a key tissue mediating a large proportion of the genetic risk for type 2 diabetes (T2D). T2D is a complex disease with multiple associated genetic risk loci identified through genome-wide association studies (GWAS) have identified >700 signals (Mahajan et al., 2018), the majority of which signals are located in non-coding regions of the genome with a presumed regulatory function (Mahajan et al., 2018). Gene discovery efforts for monogenic forms of diabetes attest to the importance of transcription factors involved in pancreatic development for normal glucose homeostasis (De Franco, 2020). Since gene regulation is highly context-specific, we hypothesized that defects in islet cell development could emerge earlier in the cell lineage in progenitor cells and that some of the regions of chromatin more accessible during development would overlap with T2D risk loci. Therefore, a comparison between human fetal endocrine and adult islet cells would present a unique opportunity to identify signals specific to endocrine development.

We compared chromatin accessibility in cells from our 12 w fetal endocrine dataset with snATAC-Seq data from adult human islets (Chiou et al., 2021b), focusing on differences between (a) adult vs. fetal beta cells and (b) adult hormone-positive cells vs. fetal EPs. We identified 146,589 differentially accessible peaks between adult and fetal beta cells, with the majority of peaks accessible in fetal beta cells (129,937 peaks) compared to adult (16,652 peaks). Next, we investigated whether T2D-risk alleles in fine-mapped credible sets at 380 loci (Mahajan et al., 2018) were differentially accessible in fetal vs. adult beta cells. We identified 34 loci that were enriched within the differential peaks in fetal beta cells including known development-specific endocrine regulatory genes such as *NEUROG3* (**Figure 5E-5F**). We also identified loci containing genes not annotated to have a functional role in endocrine development, such as *LRFN2*, a gene involved in neurite outgrowth in the brain (Li et al., 2018) where fine-mapping has previously resolved the casual variant to a single SNP (Mahajan et al., 2018) (**Figure 5G**). A similar analysis comparing adult hormone-positive cells (11,242 differentially accessible peaks) vs. fetal EPs (98,334 differentially accessible peaks) identified significant peaks at 27 T2D loci (**Figure S5E**), including the monogenic diabetes gene *HNF1B (Raile et al., 2008)* and *WDR72*, a gene involved in endocytic vesicle trafficking mediated enamel mineralization (Katsura et al., 2014) that has no reported function in endocrine development or function (**Figure S5F-G**). These results provide a framework for identifying cell-type, developmental-stage specific T2D genetic risk loci, thus generating mechanistic insights into the regulatory mechanisms of T2D-associated SNPs in the context of human pancreas development.

### Benchmarking *In Vitro* Stem Cell-derived Endocrine Cells against *In Vivo* Human Fetal Endocrine Differentiation

Taken together, our analyses thus far permitted the construction of a cellular and transcriptional roadmap of human fetal endocrine differentiation, elucidating cellular heterogeneity, inferred lineage relationships, and candidate cell fate regulators. This presented an opportunity to benchmark *in vitro* beta cell directed differentiation protocols against human fetal development *in vivo*. Current *in vitro* protocols for generating stem cell-derived beta-like cells entail the differentiation of hPSCs through a step-wise process, first to definitive endoderm, then to primitive gut tube, then the pancreatic progenitor stage, followed by an endocrine progenitor stage, and finally terminating in insulin-expressing beta-like cells (BLCs) (**Figure 6A**) (Millman et al., 2016; Pagliuca et al., 2014; Veres et al., 2019). We performed a comparative analysis of a recently published scRNA-Seq dataset generated at high temporal resolution of hPSCs undergoing directed differentiation to beta-like cells, specifically at the endocrine progenitor (stage 5) and beta-like cell (stage 6) stages (Veres et al., 2019), against our fetal endocrine scRNA-Seq dataset.

**Figure 6:**
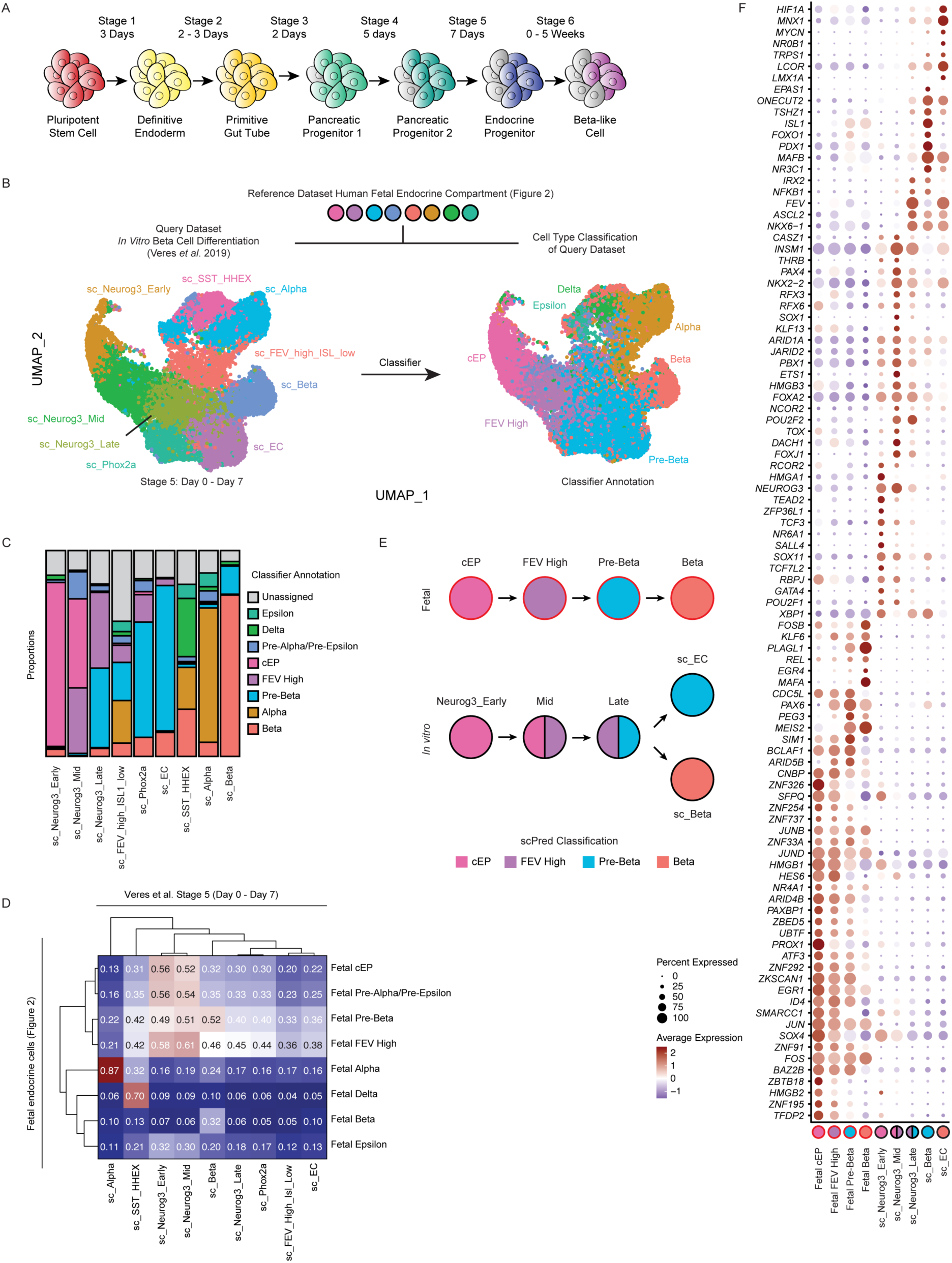
Transcriptional comparison of *in vitro* stem cell-derived endocrine cells with their endogenous *in vivo* counterparts. (A) Diagram depicting stages of *in vitro* beta cell differentiation. (B) Classifying *in vitro* stem cell-derived endocrine cells using our fetal endocrine dataset as a reference. UMAP of stem cell-derived endocrine cells generated at Stage 5 with annotation by Veres *et al*. (left) versus by cell-based classifier scPred (right) according to similarity to endogenous human fetal endocrine cells. (C) Proportions of each cluster from the Veres *et al*. dataset that are either unassigned (gray) or annotated as corresponding to a fetal endocrine cell type. (D) Heatmap depicting the Pearson correlation between *in vitro* stem cell-derived Stage 5 cells and fetal endocrine cells *in vivo* based on all shared genes. (E) Model showing inferred lineage relationships among fetal vs. *in vitro* stem cell-derived endocrine progenitor cells. (F) Dot plot depicting expression of transcription factors differentially expressed between *in vitro* and *in vivo* datasets.

First, we sought to determine whether the cellular populations generated *in vitro* are analogous to the populations we observed during human fetal development (**Figure 3A**). Veres *et al*. identified nine stem cell (sc)-derived endocrine populations that arise during the 7 days of culture at the endocrine progenitor stage (stage 5): *Neurog3+* progenitors (sc_Neurog3 Early, Mid, and Late), differentiated cell types (sc_Alpha, sc_Beta, and sc_SST_HHEX (Delta)), as well as three populations that presumably represent mis-differentiated cell types (sc_Enterochromaffin (sc_EC), sc_Phox2a, and sc_FEV_High_ISL1_Low) (**Figure 6B**). We utilized the supervised cell type classifier scPred (Alquicira-Hernandez et al., 2019) to train a prediction model using our human fetal endocrine scRNA-Seq dataset as the reference and then apply this classifier to the stage 5 endocrine cells (**Figure 6B**). The scPred classifier annotated the stage 5 sc_Beta and sc_Alpha populations as largely beta (78%) and alpha (65%) in identity, respectively, while the sc_SST_HHEX population was annotated as a mixture of alpha, beta, epsilon and delta cells, with the highest proportion (28%) of cells annotated as delta identity (**Figure 6C**). scPred classified the sc_Neurog3_Early cluster largely (80%) as the cEP population found in the fetal pancreas, while the sc_Neurog3_Mid cluster was classified as a mixture of mostly cEP (43%) and FEV High (32%) progenitors and sc_Neurog3_Late was largely classified as either the FEV High (37%) or Pre-Beta (39%) (**Figure 6B-6C, Figure S6A**). Similarly, Pearson correlation of all shared genes between the fetal and *in vitro* clusters confirmed that the four *in vitro* endocrine progenitors showed highest correlation with the fetal progenitors (**Figure 6D**). These results indicate that the transcriptional profiles of endocrine progenitor cell types found *in vitro* are largely similar to those of endocrine progenitor cell types found during human fetal development. The sc_EC and sc_Phox2a populations, which represent mis-differentiated cell types, were largely classified as the Pre-Beta (71% and 56%, respectively), likely due to the shared transcriptional networks found between pancreatic endocrine cells and enteroendocrine cells of the gut (Grün et al., 2015; Haber et al., 2017; Lavergne et al., 2020). The sc_FEV_High_ISL1_Low population had no clear classification, indicating that it does not appear to correspond to a pancreatic endocrine population present *in vivo* and instead likely represents an artifact of the *in vitro* culture platform. Taken together, our work demonstrates key similarities of *in vitro* derived human EPs and BLCs to normal development, as well as identifies the generation of cell types not found in normal development

When re-examining *in vitro* lineage relationships within the framework of the trained cell type classifier, we observed that beta cell differentiation *in vitro* occurs in a manner largely similar to endogenous human fetal development (**Figure 6E**). Veres *et al*. showed that stage 5 sc_Neurog3_Late endocrine cells can give rise not only to sc_Beta cells, but also to sc_EC cells, which resemble serotonin-secreting cells found in the gut (Veres et al., 2019). As the sc_EC population constitutes a large portion of the cells produced at the completion of the directed differentiation protocol (**Figure 6B, Figure S6B**), understanding transcriptional mechanisms that regulate their formation would aid in driving the progenitors at the previous endocrine progenitor stage to differentiate into beta cells over this unwanted population. To identify such mechanisms, we performed differential gene expression analysis among cell types along the beta cell lineage *in vivo* (cEP, FEV High, Pre-Beta, Beta) and *in vitro* (sc_Neurog3_Early, sc_Neurog3_Late, sc_Beta), as well as the sc_EC population (**Figure 6F**). This analysis resulted in 1,298 DEGs with at least 0.5 logFC in expression among all populations, 98 of which are TFs. TFs enriched in the sc_EC population included *MNX1*, while those enriched in the sc_Neurog3_Late population included *IRX2*, *FEV,* and *ASCL2*. The fetal Pre-Beta and FEV High populations showed higher expression levels of beta cell-related TFs such as *PAX6*, *PEG3*, and *MEIS2* (**Figure 6F**). The fetal beta population was enriched for *MAFA* and *PLAGL1*, while the sc_Beta population was enriched for expression of *ONECUT2, ISL1, FOXO1* and *MAFB*. In future work, inducing the expression of beta cell lineage genes in the *in vitro* terminal progenitor population (sc_Neurog3_Late) may reduce or prevent the formation of these undesired, mis-differentiated cell types and improve overall efficiency of BLC generation *in vitro*.

We next set out to determine the transcriptional differences between hormone-producing endocrine cells generated *in vitro* vs. *in vivo* by comparing *in vitro*-derived endocrine cells at the final stage of differentiation, stage 6, to their fetal counterparts. Data generated by Veres *et al*. on stage 6 cells were sampled weekly from the same differentiation flask (week 0 – week 5). Our fetal cell type classifier classified the sc_Beta and sc_Alpha hormone-expressing cell types largely as their fetal counterparts, while the sc_Neurog3 and sc_EC populations were again classified as fetal endocrine progenitor populations cEP (sc_Neurog3) and FEV High/Pre-Beta (sc_EC) (**Figure S6B**). *In vitro* beta cells sampled at the beginning of stage 6 (weeks 0 and 1) were correlated most highly with the fetal Pre-Beta population as opposed to fetal Beta cells, confirming that early stage 6 beta cells more closely resemble progenitors than differentiated beta cells and that maturation from a progenitor to a more fetal state indeed occurs over time in culture (**Figure S6C**). Differential gene expression analysis between *in vitro* and fetal beta cells revealed enrichment of genes such as *MEG3*, *PLAGL1*, *INS*, and *MAFA* in the fetal beta cell population compared to its *in vitro* counterpart, with ISH staining against *MEG3* (You et al., 2016) further confirming this observation (**Figure S6D-S6E**). Although *in vitro* beta cells showed higher expression of beta cell maturation genes *HOPX* and *IAPP* (Augsornworawat et al., 2020; Hrvatin et al., 2014), they also showed higher expression of beta cell progenitor markers such as *FEV* and *PAX4* (Byrnes et al., 2018), as well as neuronal-associated genes such as *DDC* (**Figure S6D)**. Thus, *in vitro*-derived beta cells more closely resemble fetal beta cell progenitors in the first 3 weeks of stage 6 culture before gradually more closely resembling fetal beta cells, expanding upon a previous report stating that they are more fetal-like than adult (Hrvatin et al., 2014). These results highlight the value of our dataset comprising intermediate progenitor populations at single-cell resolution in enabling the more precise mapping of *in vitro*-derived populations to their endogenous counterparts.

Next, we compared *in vitro*-derived alpha cells to their fetal counterparts and found high concordance between the two populations, irrespective of the week in culture (correlation coefficient > 0.88 at all timepoints) (**Figure S6C**). Differential gene expression analysis between the two alpha cell populations again revealed the enrichment of neuronal associated genes in the *in vitro* condition, but we observed similar expression of key alpha cell-related genes such as *ARX, IRX2, GCG, TTR* and *FOXA2* between the *in vitro* and alpha fetal cells (**Figure S6F**). These results suggest that *in vitro*-derived alpha cells closely resemble fetal alpha cells in expression of key alpha cell fate regulators, despite the fact that the differentiation protocol was optimized for the generation of beta cells (Pagliuca et al., 2014; Veres et al., 2019).

### A Functional Role for *FEV* in Regulating Human Endocrine Differentiation

Our analysis of the developing human endocrine pancreas identified *FEV* as an enriched marker of two of the EP populations within the beta cell lineage (FEV High, Pre-Beta) (**Figure 4A, Figure S3C**). Previous genetic ablation studies in the developing murine pancreas demonstrated that global knockout of the transcription factor *Fev* leads to reduced insulin production and secretion, as well as impaired glucose tolerance in adult mice (Ohta et al., 2011). The role of *FEV* in regulating human pancreas endocrine development, however, is still unknown. We set out to interrogate whether *FEV* is simply a marker of pre-beta cell populations or it itself has a functional role in endocrine development in human cells. To evaluate whether our *in vitro* stem cell differentiation platform could serve as a model for assessing *FEV* function in human endocrine development, we first asked whether the dynamics of *FEV* expression *in vitro* recapitulated those *in vivo*. *FEV* mRNA expression was first detected during the pancreatic endocrine progenitor stage (stage 4) and peaked during the endocrine progenitor stage (stage 5) **(Figure S7A)** during the differentiation. As in human fetal beta cell differentiation, *FEV* expression co-localized with *NEUROG3* at the endocrine progenitor stage **(Figure S7B).** These results indicated that FEV was expressed in human endocrine progenitor cells *in vitro* and that the stem cell differentiation platform could be used to evaluate the function of *FEV* in human EPs.

To determine the consequences of loss of *FEV* during differentiation to the beta cell lineage, we established a clonally-derived *FEV* knockout (FEV-KO) hESC line using the CRISPR-Cas9 system (**Figure 7A**) and subjected this line, alongside a WT (non-edited) control line, to directed differentiation to the beta cell lineage. As measured by flow cytometry, KO of *FEV* did not lead to any significant change in the maintenance of pluripotency as judged by staining for NANOG/OCT3/4 or in the efficiency of generating target cells at stages 1 through 5 of the differentiation, including the proportion of SOX17+/FOXA2+ cells at the end of stage 1, of PDX1+ cells at the end of stage 3, or of PDX1+/NKX6.1+ cells at the end of stage 4 or 5 (**data not shown**). However, early in the beta cell stage (stage 6, day 4), flow cytometric analysis revealed a significant reduction in the number of BLCs as measured by co-expression of both C-PEPTIDE (C-PEP, a proxy for INS) and NKX6.1, a key transcription factor of beta cells (7.9 ± 2.3%; Mean ± SEM) in the FEV-KO group vs. (22.7 ± 8.5%; Mean ± SEM) in the WT group (**Figure 7B, Figure S7C**). These results suggest that *FEV* expression is not required for pancreatic specification or induction to the endocrine lineage, but plays a critical role later in beta cell fate specification.

**Figure 7:**
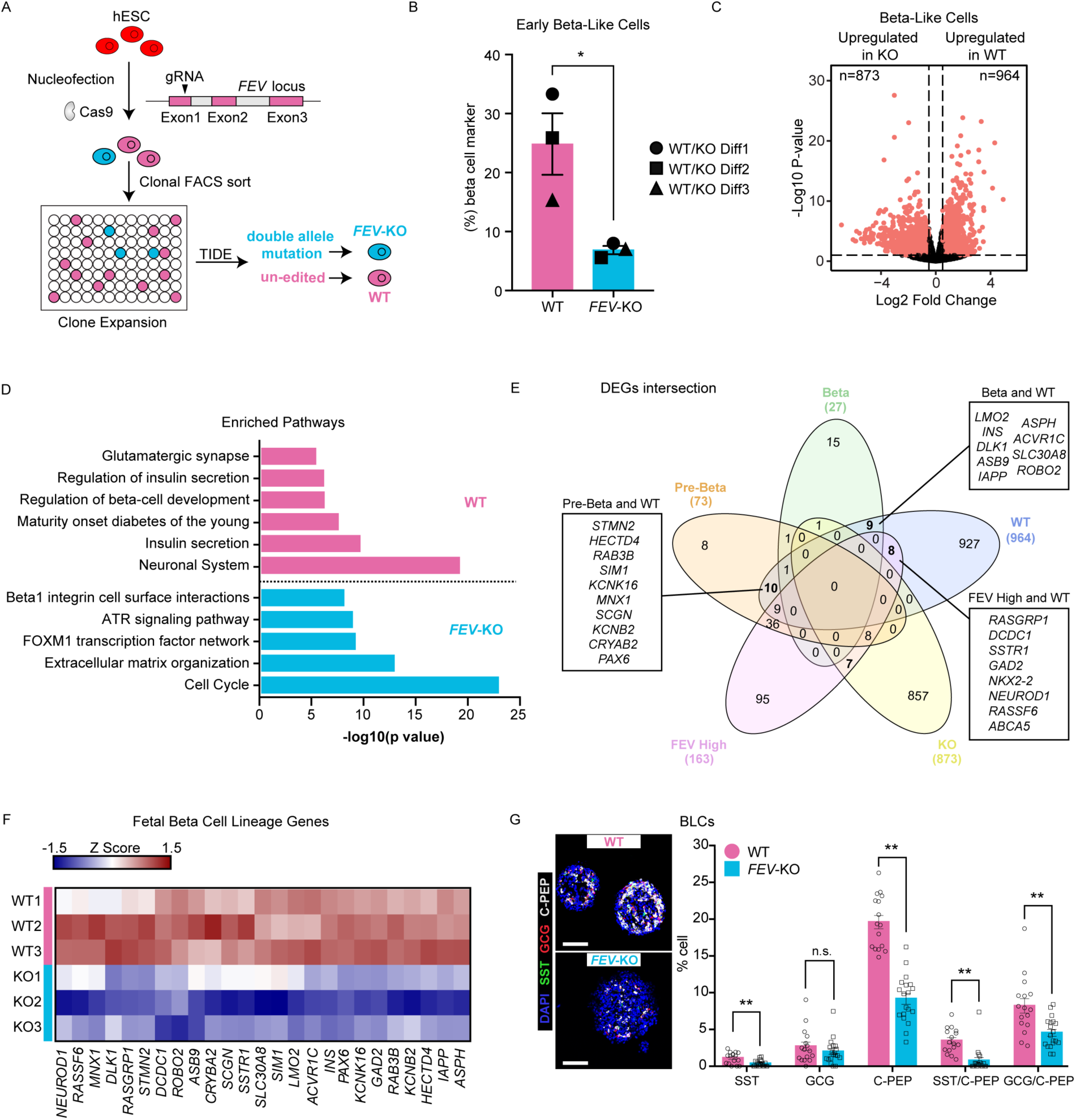
Loss of *FEV* diminishes differentiation of human stem cells to pancreatic endocrine cells *in vitro*. (A) Schematic for the generation of a *FEV* knockout (KO) human embryonic stem cell (hESC) line using CRISPR-Cas9 mediated gene editing. A WT (non-edited) line was used as a control. (B) Quantification of flow cytometry data from three independent paired differentiations of FEV-KO vs. un-edited WT control hESC towards the early beta-like cell stage (Stage 6, day 4). Efficiency of generating beta cells was quantified using staining for C-PEPTIDE and NKX6.1. Data presented as mean ± SEM (n = 3 independent batches of paired differentiation). *, p-value < 0.05, paired t-test. (C) Volcano plot depicting the genes differentially expressed between the FEV-KO vs. WT beta-like cells (BLCs) at Stage 6, day 10 of the directed differentiation, as assessed by bulk RNA-Sequencing. Red dots depict genes with a log 2-fold change (FC) of at least 0.5. (D) Pathway analysis of the genes differentially expressed between FEV-KO vs. WT cells at Stage 6, day 10. (E) Five-way Venn diagram showing intersection of differentially-expressed genes (DEGs) (defined as log2 FC > 0.5) specific to any one of the fetal beta lineage cell populations (*i.e.,* FEV High, Pre-Beta, and Beta) along with DEGs identified in a comparison of WT vs. KO BLCs. (F) Heatmap depicting the expression levels of representative fetal beta lineage genes in paired differentiations of FEV-KO and WT cells. (G) Representative staining of WT and FEV-KO BLCs (Stage 6, day 12) for SST (green), GCG (red), C-PEP (gray), and DAPI (blue) (left panels). Scale bars: 100 um. Right panel: Quantification of aggregate immunofluorescence staining data across two batches of differentiation (N=1770 WT cells, N=2300 FEV-KO cells). Each dot represents the cell ratio quantified in one cluster, graphs are represented as mean ± SEM; n.s., not significant; ** P < 0.01; unpaired t-test.

To understand the transcriptional changes upon KO of *FEV*, we performed bulk RNA-Sequencing (RNA-Seq) on FEV-KO and WT cells from three independent, paired differentiations at the BLC stage (Stage 6, day 10). As expected, we observed very few genes (< 10 genes per timepoint) differentially expressed between WT and KO conditions at the end of stages 1 and 4, time points before peak *FEV* expression (**data not shown**). In contrast, at Stage 6, day 10, differential gene expression analysis between the WT and KO cells at the BLC stage resulted in 1,837 genes with at least 0.5 logFC change in expression (964 genes in WT, 873 genes in KO) (**Figure 7C**). Pathway analysis of the DEGs identified enrichment of pathways related to beta cell function, such as insulin secretion and calcium signaling in the WT condition, while the KO condition displayed enriched pathways such as cell cycle and extracellular matrix organization (**Figure 7D**). Among the DEGs were known direct targets of *Fev* in the murine pancreas and brain, including *SLC6A4, GCK, LMX1B* and *DDC*, confirming the efficacy of our CRISPR-generated KO line (**Figure S7D**) (Ohta et al., 2011; Wyler et al., 2016).

To determine whether KO of *FEV* had affected the expression of regulators of beta cell fate, we analyzed the intersection of genes identified by bulk RNA-Seq as being enriched in either the WT or KO beta-like cells with genes identified by scRNA-Seq as being differentially expressed across the beta lineage populations during development. This analysis revealed that *FEV* ablation resulted in down-regulation of fate regulators enriched in FEV high and Pre-Beta cells, including *NKX2.2*, *NEUROD1*, and *PAX6* (**Figure 7E**), as well as regulators of beta cell maturation and function such as *INS, MNX1,* and *IAPP* (**Figure 7F**). In addition, the KO of *FEV* also led to the reduction in alpha, delta- and epsilon-associated genes such as *GCG, IRX1, TTR,* and *SST* (**Figure S7G-S7H**). To verify these findings, we performed IF staining of hormone markers on the WT and FEV-KO cell clusters at the beta-like cell stage (stage 6, day 12) and observed a significant decrease in the proportion of SST+ or C-PEP+ hormonal cells in KO versus WT cells. The ratios of SST^+^C-PEP^+^ and GCG^+^C-PEP^+^ double positive bi-hormonal cells were also significantly reduced in the KO condition (**Figure 7G**). Taken together, these data indicate that *FEV is* not simply a marker of pre-beta EP cells but indeed also plays a role in regulating endocrine cell specification in the developing human pancreas.

## DISCUSSION

In this study, we have comprehensively characterized the transcriptome of human fetal pancreas at six developmental time points ranging from 8 w to 20 w at single-cell resolution. We have identified previously unappreciated levels of heterogeneity within the various pancreatic cell types, including 15 mesenchymal, 8 exocrine, 8 endocrine, 8 endothelial, 13 neuronal, 31 immune, and 20 proliferating clusters. This resource provides a cellular and gene regulatory roadmap of early human fetal pancreas organogenesis and lays the groundwork for the interrogation of the functional significance of each of these cell types. We confirmed the presence of representative subpopulations in the endothelial and mesenchymal lineages by *in situ* hybridization in fetal tissue. Future studies are warranted to validate other newly described populations and assess their presence or absence at earlier or later developmental time points not covered by this study.

By computational inference we have observed active cell-cell communication between Broad Groups in the developing human pancreas. In particular, analysis using CellChat inferred NOTCH-JAG and PDGF-PDGFR as potentially functional ligand-receptor pairs within the endothelial lineage and the mesenchymal lineage, respectively. By systematically interrogating the signaling interactions between the Endocrine and other Broad Groups, we have also identified ACTIVIN as a ligand that is expressed solely by the mature VSM cells and is predicted to act specifically on the endocrine progenitor populations. Given these data and previous evidence that non-epithelial cells such as endothelial, neuronal, and mesenchymal play important roles in guiding the development of the murine pancreatic epithelium (Attali et al., 2007; Bhushan et al., 2001; Borden et al., 2013; Cozzitorto et al., 2020; Lammert et al., 2001), the signaling pathways identified here warrant future functional confirmation by methods such as genetic or small molecule-mediated loss- and gain-of-function experiments in human pancreatic tissue *ex vivo* or using hPSC-derived pancreatic cells *in vitro*.

Our detailed investigation of the human fetal endocrine compartment identified four novel progenitor populations that each express unique marker genes. Multiplexed *ISH*/IF staining confirmed the existence of these populations across multiple developmental time points and in independent biological samples *in vivo*. Recently, an independent study identified endocrine subpopulations, several of which (termed EP2, EP3, EP4) appear analogous to the ones we have described here, providing additional evidence of their existence *in vivo (Yu et al., 2021)*. In addition, the high concordance among our *in silico* replicates gave us high confidence that these represent bona fide cellular populations in tissue (**Figure S1C**). The CellFindR clustering algorithm was critical in the discovery of these EP populations, as initial analyses utilizing standard clustering methods failed to distinguish these populations from one another and instead annotated all four as belonging to a single EP cluster. Of note, no evidence of heterogeneity was found within each differentiated hormone-producing endocrine cell type, a topic that is debated in the adult human pancreas (Baron et al., 2016; Blodgett et al., 2016; Mawla and Huising, 2019). Given the sensitivity of the clustering algorithm CellFindR in identifying cellular heterogeneity, it is likely that these heterogeneous populations do not exist in the fetal endocrine pancreas during the developmental time points covered by this study, although it is possible that they were too rare to detect or that they only arise later in development. Additionally, by adopting a variety of computational methods, we have inferred the gene regulatory networks and differentially expressed genes among endocrine populations. These insights serve to significantly improve our understanding of human pancreatic endocrine development at the cellular and molecular levels.

The endocrine cell lineage predictions constructed for human cells in this study provide a contrasting account of endocrine differentiation when compared to murine development. Unlike in the mouse pancreas, where multipotent intermediate progenitors give rise to all endocrine cell types (Bastidas-Ponce et al., 2019; Byrnes et al., 2018; Yu et al., 2019b), our data presented here predict that three of the four human EP populations act as fate-committed progenitors that are either uni- or bi-potent with respect to a specific hormone-expressing cell type. This differentiation potency is reflected in the genes expressed in these populations, as transcriptional analysis revealed gradual increase of expression of differentiated endocrine cell related genes as the EPs began to become more fate restricted. The alpha and beta lineage predictions presented here are consistent with a recent study that utilized mitochondrial genome variants within adult alpha and beta cells as endogenous lineage tracing markers at single-cell resolution and concluded that human alpha and beta cells arise from separate progenitor populations (Lin et al., 2021). Prediction of the delta lineage presented here, however, is less clear. In particular, the FEV High and Delta clusters were not connected in the UMAP and may have been “forced” into the same lineage by the computational reconstruction **(Figure 3A-3B)**. Of note, our lineage analysis of human endocrine differentiation differs from a recently published study using single-cell sequencing and lineage reconstruction of fetal endocrine cells (Yu et al., 2021). Future studies at early developmental time points might resolve this issue by increasing the cell number of captured endocrine progenitors, increasing the chances of detecting potentially rare and/or transient, pre-delta progenitor cells. In the future, applying state-of-the-art methods for lineage barcoding to human pancreas tissue *ex vivo* could represent an exciting approach for experimental validation of predictions generated *in silico*.

The lineage predictions based on scRNA-seq data are consistent with those generated by snATAC-seq analysis, which has validated the existence of each of the four novel EP subtypes by an orthogonal method. Furthermore, the multi-omic approach described in this study represents a unique resource for identifying candidate regulators of endocrine cell fate specification in the human fetal pancreas. By overlapping the “active” regulons identified by SCENIC analysis and motif enrichment/RNA expression correlation analysis (“positive” TFs), we have gained a clearer picture of the TFs most likely to be relevant to endocrine differentiation and maturation. For instance, the TCF family of TFs (*TCF4*, *TCF12*, *TCF3)*, which are known regulators of endocrine differentiation (Jacquemin et al., 2000), were identified by both analyses as TFs that are potentially mediating gene expression and chromatin accessibility in endocrine cells. Additionally, these analyses also identified TFs with unknown endocrine development function, such as *CUX1 and POU2F2*. As such, the data presented here should serve as a resource for the field with broad utility in identifying cell fate regulators in future studies.

Identification of the effector transcripts or genes through which disease associated variants influence risk is a critical first step towards biological inference and thus clinical translation. The context-specificity of gene regulation presents an additional challenge. Single-cell resolution of both gene expression and chromatin accessibility in human adult islets has recently demonstrated the importance of cell state, cell type and the potential of co-accessibility analysis between promoters and cis-regulatory elements to identify effector genes at T2D-risk loci (Chiou et al., 2019, 2021b; Rai et al., 2020). Here, we have extended this to include the influence of developmental stage by performing a comparative analysis of human adult vs. fetal endocrine cells to uncover enrichment of T2D genetic risk loci, permitting the assessment of regions of DNA harboring T2D-risk alleles that are accessible during development and may therefore affect the expression of developmental genes. As expected, our analysis identified fetal enrichment of risk loci of known regulators of endocrine differentiation such as *NEUROG3*. We also identified, however, fetal enrichment in genes with no known function in endocrine development, such as *LRFN2*. One intriguing observation from our analysis is the potential for further context specificity on gene regulation at the *PROX1* locus. There are two independent signals at this locus: the first has been fine mapped to a single variant (rs340874), and the second has two SNPs (rs79687284 and rs17712208) in the credible set (Mahajan et al., 2020). An evaluation of these variants on transcriptional activity in both human HepG2 hepatocytes and EndoC-βH1 beta cell models using *in vitro* reporter assays demonstrated effects in both liver and beta cells for rs340874; at the second signal, however, only one of the two variants (rs17712208) influenced activity in beta cells and neither in HepG2 cells (Mahajan et al., 2020). Our data now raise the intriguing possibility that the rs79687284 variant could alter activity earlier in development, thus expanding the complexity of the regulatory impact at this locus. These analyses therefore provide a framework for the identification of development-specific disease risk loci and a rich opportunity for further study of their function in islet biology.

Despite tremendous progress in recent decades in devising methods to generate beta-like cells *in vitro* from hPSCs, these protocols still suffer from the generation of unwanted cell types. A previous study aimed to assign hPSC-derived endocrine cell identification by referring to adult islets cells, but was constrained by the absence of endocrine progenitors in adult tissue (Krentz et al., 2018). By performing computational comparison of endogenous *in vivo* vs. *in vitro* endocrine development, we observed that the EP cell types made *in vitro* are similar to those present *in vivo*. That said, the generation of mis-differentiated cell types such as the stem cell-derived enterochromaffin (sc_EC) population demonstrates that there remains significant room for improvement of the differentiation protocol with respect to purity and efficiency. Given the similarities between pancreas and intestinal endocrine development, the generation of the enterochromaffin population is likely due to the mis-expression of key genes that then tips the balance towards an enteroendocrine fate. Future work will focus on modulating the expression, ideally in a temporally constrained fashion, of key genes that are currently aberrantly expressed in order to generate more pure and functionally mature beta cell populations. Additionally, our data have important implications for the generation of *in vitro-*derived islet cells, as our *in vivo* developmental roadmap can now be used for the refined production of non-beta endocrine cells, including human alpha (Peterson et al., 2020; Rezania et al., 2011) and delta cells.

Lastly, the results of our *FEV* gene ablation study demonstrate how the generation of a detailed *in vivo* roadmap of endocrine differentiation can successfully be combined with *in vitro* genome editing techniques to discover important regulators of human endocrine development.

We verified that *FEV* not only marks beta cell progenitors in the developing fetal pancreas, but itself also plays a functional role in human endocrine differentiation *in vitro*. Further investigation of *FEV* through the use of TF binding studies will provide insight as to how it regulates endocrine differentiation or function. Applying this approach to other genes of interest is a promising approach for understanding additional, uncharacterized regulators of human endocrine differentiation.

In summary, we provide here a comprehensive, single-cell, multi-omic roadmap of human fetal pancreatic endocrine development. This study represents a critical step towards generating bona fide beta cells *in vitro* for therapeutic use.

### Limitations of Study

Due to constraints on access to human fetal tissue, our study is necessarily limited to a window of pancreatic development after which endocrine development has already been initiated, and before full maturation of fetal hormone-expressing cell types has occurred. Future work utilizing techniques such as laser capture microdissection or other methods may permit a bridging of the gap between the latest timepoints possible in our study (20 w) and other work that has performed single-cell transcriptional analysis of adult human pancreas tissue. As with any computational methods that rely on inference of lineage relationships, our work on lineage reconstruction of fetal endocrine development should be interpreted with caution, as classical experimental lineage tracing techniques are not possible in human tissue.

The snATAC-Seq analysis in this study was performed on human tissue from a single timepoint and, despite enrichment of epithelial cells, suffers from low cell numbers within several endocrine subgroups, which prevents the robust interpretation of lineage relationships based on chromatin accessibility. Performing snATAC-Seq on additional samples and timepoints would provide a better understanding of the epigenetic mechanisms regulating human pancreatic endocrine development.

## Methods

### RESOURCE AVAILABILITY

#### Lead Contact

Further information and requests for resources and reagents should be directed to the Lead Contact, Julie Sneddon.

#### Materials Availability

All unique/stable reagents generated in this study are available from the Lead Contact with a completed Materials Transfer Agreement.

#### Data and Code Availability

Scripts used in this study will be made available at GitHub.

Raw single-cell sequencing data of human fetal pancreas samples will be made available in dbGaP, and raw and processed data of the FEV WT and KO bulk RNA sequencing will be made available in GEO.

### EXPERIMENTAL SUBJECT DETAILS

Informed consent was obtained for all human tissue collection, and protocols were approved by the Human Research Protection Program Committee at UCSF. Human fetal dorsal pancreas tissue was obtained from post-mortem fetuses at 8 to 20 weeks post conception (w) through two sources: University of Washington Birth Defects Research Laboratory (BDRL) and Advanced Bioscience Resources, Inc. (ABR). Identifiers were maintained at the source only, and the investigators received only de-identified specimens. After isolation, tissue was shipped overnight (O/N) on ice in RPMI medium. A portion of tissue was fixed in 4% paraformaldehyde O/N at 4 °C, washed three times with 1 x phosphate-buffered saline (PBS), and cryopreserved in 30% sucrose solution at 4 °C for O/N in preparation for embedding in optimal cutting temperature (OCT) compound. Sections measuring 10 um in thickness were cut using a cryostat and stored at -80°C for immunofluorescence staining or *in situ* hybridization, as described below.

Adult human islets were isolated from cadaveric donor tissue by the UCSF Islet Production Core with permission from the UCSF ethical committee. Consented cadaver donor pancreata were provided by the nationally recognized organization UNOS via local organ procurement agencies. The identifiers were maintained only at the source, and the investigators received de-identified specimens.

### METHODS DETAILS

#### Processing of pancreas tissue scRNA-Seq and snATAC-Seq

To isolate cells for single-cell RNA-Sequencing, human fetal pancreas tissue was minced with scalpels and transferred to dissociation buffer containing Liberase TM and 0.1 mg/mL Dnase I for 30-55 minutes at 37°C on a Thermomixer at 1000 rpm. Enzyme was quenched with 1X HBSS containing 5mM EDTA and 10% FBS. The resulting cell suspension was filtered through a 30 um strainer. All tissues were subject to removal of red blood cells (RBCs) using immunomagnetic negative selection with the EasySep RBC Depletion kit (STEMCELL Technologies, 18170). 12 w samples were further subjected to EasySep™ Human EpCAM Positive Selection Kit II (STEMCELL Technologies, 18356) to positively select for epithelial cells. Tissues at 19 w and 20 w were subjected to EasySep™ Human CD45 Depletion Kit II (STEMCELL Technologies, 18259) to remove CD45+ immune cells. Cell viability was measured for all samples using a MoxiFlow (Orflo) to confirm greater than 90% viability.

To isolate nuclei for single-nuclei ATAC-Sequencing, 12 w human fetal pancreatic tissue was placed in a dissociation buffer containing Liberase TM and 0.1 mg/mL Dnase I at 37 °C. Dissociated cells were filtered through a 30 um strainer and further enriched for EpCAM+ epithelial cells using the EasySep™ Human EpCAM Positive Selection Kit II. Nuclei from EpCAM+ cells were isolated following 10x Genomics protocol CG000169, Rev D. In brief, EpCAM+ cells were resuspended in PBS + 0.04% BSA and centrifuged at 1000 rpm and 4°C for 5 min. Chilled Lysis Buffer was added to the cell pellet, which was then incubated on ice for 3.5 min. After lysis, chilled Wash Buffer was added, cells were centrifuged at 1200 rpm, and isolated nuclei were suspended in 1X Nuclei Buffer. After isolation, nuclei were manually counted with a hemocytometer; quality was assessed under a 63x bright field microscope to ensure that the periphery of isolated nuclei appeared smooth.

#### Single-cell capture and sequencing

For scRNA-Seq of human fetal tissue, we utilized the Chromium Single Cell 3’ Reagent Version 3.1 Kit (10x Genomics). For non-enriched human fetal samples, we loaded 25,000 cells each onto two lanes of the 10x chip, resulting in a total of 50,000 cells loaded per sample. For enriched human fetal samples, including EpCAM+ cells from two 12 w samples, EpCAM+ cells and CD45-cells from one 19 w sample, and CD45-cells from one 20 w sample, 25,000 cells from each enrichment condition were loaded onto a single lane of the 10x chip. Gel Bead-In EMulsions (GEMs) were generated and subjected to reverse transcription for RNA barcoding before cleanup and cDNA amplification. Libraries were then prepared with the Chromium Single Cell 3’ Reagent Version 3.1 Kit according to the manufacturer’s instructions. Each resulting library was sequenced on the Novaseq 6000 platform (Illumina) with the following parameters: Read 1 – 28 cycles, Index 1 i7 – 8 cycles, Index 2 i5 – 0 cycles, Read 2 – 91 cycles.

For snATAC-Seq of 12w human fetal tissue, we utilized Chromium Next GEM Single Cell ATAC Library & Gel Bead Kit v1.1 (10x Genomics). 7,166 nuclei were loaded onto one lane of a 10x chip. Transposition, GEM generation and barcoding, cleanup and library construction were performed according to the manufacturer’s protocol. The library was then sequenced on the Novaseq 6000 platform (Illumina) with the following parameters: Read1-50 cycles, Index1 – 8 cycles, Index 2 – 16 cycles, Read 2 – 49 cycles.

#### Human fetal single-cell RNA-Sequencing analysis

To assemble the transcriptomic profiles of individual cells, we utilized CellRanger versions v3.0-4.0 with default settings to demultiplex, aligned reads to the human genome (GRCh38, supplied by 10x Genomics), and quantified unique molecular identifiers (UMIs). The resulting gene-barcode matrices were then analyzed and aggregated with the R package Seurat v3.1.2 (Stuart et al., 2019). Each sample was subjected to filtering to exclude cells expressing fewer than 200 genes and genes expressed in fewer than three cells. Technical replicates (two 10x lanes of the same biological sample) were merged using the MergeSeurat() function. High-quality cells were retained by filtering on the number of expressed genes and mitochondrial content. Each sample was normalized with NormalizeData(), and variable genes were identified with the FindVariableFeatures() function using 2,000 genes and the “vst” selection method. Integration anchors were found across all samples with the FindIntegrationAnchors() with 30 principal components and 2,000 genes. The samples were then integrated using the IntegrateData() function. The data was then scaled with ScaleData() function and principal component analysis (PCA) was performed, with 30 principal components selected based on the ElbowPlot(). Dimensionality reduction and initial clustering was performed with the FindNeighbors(), FindClusters() and RunUMAP() functions using 30 principal components and a resolution parameter of 2.0. The resulting Louvain clusters were then manually annotated into “Broad Groups” of known biological cell types using canonical marker genes. Cluster 17 from the initial clustering was removed, as it had no distinguishable marker genes and expressed low levels of features.

#### Cell clustering with CellFindR

To further sub-cluster the Broad Groups, we applied a novel clustering package, CellFindR (https://github.com/kevyu27/CellFindR) (Yu et al., 2019a), to each Big Group individually. Each Broad Group was subsetted, PCA and UMAP were recalculated, and the top level resolution was found with the res() function. Iterative sub-clustering was performed on each top level cluster with the sub_clustering() function. Clusters that were deemed non-biological (i.e. *COL3A1*+/hormone+ doublets) were manually removed from the endocrine data set, including a sub-cluster of *INS/CELA3A-*high cells that displayed a low number of features and counts, likely representing empty droplets containing ambient RNAs known to contaminate scRNA-Seq datasets of the pancreas (Nieuwenhuis et al., 2020).

#### Single cell differential gene expression analysis

Marker genes were identified with Seurat’s FindAllMarkers() function and visualized with FeaturePlot(), VlnPlot() and DoHeatmap() functions. Pairwise volcano plots were created by utilizing the FindMarkers() function and plotting the results as a volcano plot from the EnhancedVolcano package (https://bioconductor.org/packages/release/bioc/html/EnhancedVolcano.html). Transcription factors were identified through comparison to AnimalTFDB3.0 database (Hu et al., 2019).

#### Cell-cell communication analysis

To infer cell-cell communication within our human fetal pancreas dataset, we utilized the R package CellChat (https://github.com/sqjin/CellChat). For each analysis, we used normalized counts and cell-type specific labeling as input. We then followed the ‘Inference and analysis of cell-cell communication using CellChat’ vignette.

#### Pseudotemporal ordering analysis of scRNA-Seq data

For trajectory and pseudotime analyses, we utilized the R package Slingshot (https://github.com/kstreet13/slingshot) (Street et al., 2018). Seurat-based UMAP dimensional reduction and CellFindR clustering were used as input for the merged human fetal endocrine analysis. Lineage reconstruction was performed with the slingshot() function, with the cEP population (Cluster 3.1.0) designated as the beginning of pseudotime.

#### Pathway analysis

Pathway analysis and calculation of associated p values were performed using the ConsensusPathDB overrepresentation analysis for pathway based sets category (http://cpdb.molgen.mpg.de).

#### Gene regulatory network analysis

For gene regulatory network (GRN) analysis, we utilized the R package SCENIC (Single-Cell rEgulatory Network Inference and Clustering; https://github.com/aertslab/SCENIC) (Aibar et al., 2017) and the PYTHON package GRNBoost (https://github.com/aertslab/GRNBoost) (Aibar et al., 2017) The 500bp-upstream and tss-centered-10kb human RcisTarget database were selected for analysis, and RNA counts from the merged human fetal endocrine pancreas dataset were utilized as input. Genes were retained only if they had at least 6 UMI counts, were detected in at least 1% of cells, and were available in the RcisTarget database. To distinguish potential activation from repression, we calculated the correlation with the runCorrelation() function. The resulting filtered expression matrix and identified transcription factors were analyzed with GRNBoost for GRN inference. Finally, the GRNs were built and scored with the SCENIC functions runSCENIC_1_coexNetwork2modules(), runSCENIC_2_createRegulons(), and runSCENIC_3_scoreCells. The resulting scaled regulon activity AUC scores were displayed via heatmaps with the R package ComplexHeatmaps function.

#### Initial snATAC-Seq analysis

To assemble the chromatin profiles of individual cells, we utilized Cell Ranger ATAC v1.1 with default settings to demultiplex, align reads to the human genome (using the pre-built GRCh38 human genome supplied by 10x Genomics), and generate single-cell accessibility counts. The resulting files were then analyzed using the R package ArchR v0.9.5 (Granja et al., 2020). Arrow files were created from the fragment file from the CellRanger output with the createArrowFiles() function and an ArchR project created with the ArchRProject() function. Doublets were filtered out using the addDoubletScores() and filterDoublets() functions, resulting in a dataset comprising 6,010 nuclei. Dimensional reduction was calculated with the addIterativeLSI() function using the following settings: iterations = 2, resolution = 0.5, sampleCells = 2500, n.start = 10, varFeatures = 25000, dimsToUse = 1:30. Clustering was performed with addClusters() with method = Seurat and resolution = 0.1. UMAP was calculated with addUMAP() and clusters were annotated as “Broad Groups” based on marker gene expression (see above in Initial single-cell RNA-Sequencing analysis).

The endocrine cluster was then subsetted and dimensional reduction was calculated with the addIterativeLSI() function with the following settings: iterations = 2, resolution = 0.3, sampleCells = 1500, n.start = 10, varFeatures = 25000, dimsToUse = 2:10. UMAP was calculated with addUMAP() with default settings. Unconstrained integration with scRNA-Seq data was then performed using the final scRNA-Seq endocrine dataset (Fig. 2a) as input with the addIntegrationMatrix() function, transferring cluster labels and a pseudo-RNA-Seq profile. To maintain the robustness of the analysis, the endocrine progenitor (EP) clusters were collapsed into a single EP cluster, resulting in a final dataset comprising 5 clusters and 1,754 cells. **snATAC-Seq peak calling**

To call peaks in the snATAC-Seq dataset, group coverages were added to the ArchR project with addGroupCoverages() and peaks called with Macs2 with the addReproduciblePeakSet() function with a cutoff of 0.1. The peak set was then added to the project with addPeakMatrix(). Marker peaks were identified with the getMarkerFeatures() function using the PeakMatrix and peaks with a false discovery rate (FDR) less than or equal to 0.1 and log2 fold-change of 0.5 or greater were visualized with markerHeatmap().

#### Motif analysis

To perform motif enrichment in regions of open chromatin, we first added motif annotations with the addMotifAnnotations() function, using “cisbp” as the motif set. To visualize motif deviation at single-cell resolution, we utilized ChromVAR (Schep et al., 2017) within the ArchR package. Background peaks were added with addBgdPeaks() and per-cell deviations calculated with addDeviationsMatrix(). Deviation scores were then visualized with the plotEmbedding() function.

To calculate “positive” transcription factors (TFs showing both motif enrichment and RNA expression), deviant motifs were first accessed with seGroupMotif(). The motif deviation scores and RNA expression profiles were then correlated with the correlateMatrices() function and filtered based on a threshold of correlation greater than 0.5, adjusted p-value less than 0.01 and a maximum inter-cluster difference in deviation z-score that is in the top quartile.

#### Pseudotemporal ordering analysis of snATAC-Seq data

To order cells in pseudotime, we first increased the clustering resolution, as ArchR does not allow for ordering of trajectories with fewer than 3 clusters. Clustering was performed with the addClusters() function with a resolution setting of 1.0. Alpha, Beta, Delta and Epsilon trajectories were then manually chosen by clusters; the cells along those trajectories were then ordered by pseudotime with the addTrajectory() function. The trajectories were then visualized with plotTrajectory().

To assess positive transcription factors across pseudotime, we accessed motif enrichment and RNA expression across pseudotime with getTrajectory(), using the motif matrix and gene integration matrix, respectively. We then correlated these trajectories with cutoffs of correlation = 0.5 and variance quartile cutoff = 0.8 for both the motif matrix and gene integration matrix.

#### T2D GWAS enrichment analysis

Datasets for adult islet samples were accessed from GEO (GSE160472, Chiou et al., 2021) and analyzed together with our fetal pancreas sample. Combinatorial barcoding (CB) data were processed with the ENCODE ATAC-seq pipeline (v1.9.3) by aligning to the genome reference GRCh38. Cell barcoding information contained in the read names was added as CB tags in the bam files with a customized script. Only mapped reads with MAPQ score > 30 were retained. Cell type annotation of adult islet cells were applied from the metadata file provided on GEO. ArchR (v1.0.1) was used as a platform for the downstream analysis, including clustering, peak calling (MACS2, v2.2.7.1) and wilcoxon testing for differential peaks. T2D loci were prioritized (percentage of SNPs > 20%) based on the overlapping between significant peaks (FDR < 0.05 & abs(Log2FC) > 1) and the SNPs in the 99% genetic credible interval for 380 distinct T2D association signals ((Mahajan et al., 2018) with GRCh37 coordinates mapped to GRCh38 by LiftOver).

#### Analysis of scRNA-Seq data from *in vitro*-differentiated stem cell-derived cells

Processed counts and cell metadata were downloaded from GEO accession number GSE103412 for Stage 5 (EP stage) and stage 6 (beta cell stage) cells (Veres et al., 2019). Counts and meta data were read into R and Seurat objects were created with associated metadata. Data were normalized with NormalizeData() and variable features found with FindVariableFeatures(). Data were scaled and PCAs calculated with ScaleData() and RunPCA(), then data were clustered with FindNeighbors(), FindClusters() and data reduced with RunUMAP().

Endocrine cell types were classified with the R package scPred (Alquicira-Hernandez et al., 2019). First, the features used for classification were calculated on the human fetal CellFindR endocrine clusters with getFeatureSpace(). The classifier was then trained with trainModel(), with “mda” as the model. The classifier was then applied to the stage 5 and stage 6 datasets with ScPredict().

For *in vitro* and *in vivo* differential gene expression analysis, datasets were combined with the MergeSeurat() function and differential gene expression calculated as stated above.

#### Multiplexed *in situ* hybridization and immunofluorescence

Human fetal pancreas tissue was fixed in 4% paraformaldehyde (PFA) overnight (O/N) at 4 °C. hESC-derived clusters were fixed with 4% PFA for 20 min at room temperature (RT). Post-fixed tissues and clusters were washed three times with 1x phosphate-buffered saline (PBS), cryopreserved in 30% sucrose solution at 4 °C O/N, and embedded in optimal cutting temperature (OCT) compound. Sections measuring 10 um in thickness were cut using a cryostat and stored at -80 °C for *in situ* hybridization and immunofluorescence, as described below.

After removal from -80 °C storage and incubation at RT for 30 minutes, cryosections were washed with 1 x PBS to remove OCT, and sequentially treated with hydrogen peroxide and proteinase III. Tissues were then hybridized with probe mixes for 2 hours at 40 °C. Inventoried or customized probes were purchased from Advanced Cell Diagnostics, Inc. Probes against *SUSD2* (42673), *PRPH* (410231-C2), *FEV* (471421-C3), *NEUROG3* (050798-C4), *LMX1B* (582661), *MEG3* (400821), *ARX* (486711-C2), *SFRP1* (428381-C4), *SFRP2* (476341-C3), *CCL21* (474371-C2), *GJA5* (471431-C2), *PLVAP* (437461), and *ACKR1* (515131) were used according to the manufacturer’s instructions for the RNAScope multiplex fluorescent detection V2 kit (Advanced Cell Diagnostics, Inc, 323110). To validate probe specificity, negative control probe (DapB) and positive control probe (*POLR2A*/*PPIB*/*UBC*) were included in each experiment. Hybridization signals were amplified via sequential hybridization of amplifier AMP1, AMP2, and AMP3 and fluorophores Opal 570 (1:1500, PerkinElmer, FP1488001KT), Opal 650 (1:1500, PerkinElmer, FP1496001KT), Opal 690 (1:1500, PerkinElmer, FP1497001KT). Following signal amplification of the target probes, sections were either stained with DAPI and mounted for imaging, or continued with standard immunofluorescence (IF) procedure. For IF, sections were incubated in 1 x blocking buffer (0.1% PBST containing 5% normal donkey serum) for 1 hr at RT then stained O/N at 4° C using the following primary antibodies diluted in blocking buffer: Chromogranin A (1:200, Abcam, ab15160), Glucagon (1:200, Cell Signaling, 2760S), Insulin (1:200, DAKO, A0564), Somatostatin (1:200, Santa Cruz Biotechnology, sc-7819), SMA (1:200, Abcam, ab21027), PECAM (1:100, Dako, M0823). The next day, sections were washed in 1X PBS three times and incubated with species-specific Alexa Fluor 488-, 555-, 594-, or 647-conjugated secondary antibodies (1:500) and DAPI in the blocking buffer for 1 hour at RT. Stained slides were mounted with ProLong Gold Antifade Mountant (ThermoFisher SCIENTIFIC, P36930) and stored at 4 °C prior to imaging.

#### Image acquisition

Optical sectioning images were acquired with a Leica confocal laser scanning SP8 microscope equipped with white light sources. Z-sections were captured for each imaging area with 10 steps x 1 mm thickness.

#### Genetic engineering to generate the FEV-KO hESC line

The HUES8 human embryonic stem cell (hESC) line was grown on Matrigel (Corning, 354230)-coated tissue culture plates in mTeSR1 (STEMCELL Technologies). Media was changed to mTeSR1 + 10 μM Rock inhibitor Y-27632 for 2 hours prior to nucleofection. Cells were dissociated into a single-cell suspension using TrypLE Express (Gibco, 11588846). A FEV-KO gRNA (5’-CTGATCAACATGTACCTGCC-3’) was designed using Benchling software and purchased from Dharmacon. To carry out the nucleofection, 160uM tracrRNA and 160uM FEV-KO gRNA were mixed together to make the RNA-complex and incubated for 30 min. in a 37 °C cell culture incubator. Purified Cas9-NLS protein (QB3 UC Berkeley MacroLab) was added to the RNA-complex, gently mixed to make the RNP (ribonucleoprotein), and incubated at 37 °C. After 15 min., dissociated cells were resuspended in P3 buffer (Lonza, V4XP-3032). Cell suspension and RNP were mixed and inserted into the Lonza 4D-Nucelofector (Lonza, AAF-1002B) and nucleofected in the P3 buffer. Nucleofected cells were transferred to mTeSR1 supplemented with Rock inhibitor, then seeded onto Matrigel-coated T75 tissue culture flasks (ThermoFisher Scientific, 159910).

#### Validation of FEV-KO and -WT hESC lines

Cells were sorted with FACS and clonally plated onto Matrigel-coated 96 well plates and grown in mTeSR1 supplemented with Rock inhibitor Y-27632. Clonal colonies were hand-picked under a colony-picking microscope under sterile conditions and each colony was transferred into one well of a 96-well plate, then successively passed onto larger plate formats. To determine the efficiency of genomic editing of each colony, genomic DNA from each colony was harvested with QuickExtract DNA Extraction (Lucigen, QE09050) and then used for PCR amplification. The following forward and reverse primers targeting the FEV-KO editing site were used to produce a 491-bp amplicon: 5’-CCGTCTTCTCCTCCTTGTCACC3’ and 5’-CTCGGCCACAGAGTACTCCAC-3’. PCR polymerase capable of handling GC-rich amplicons was used (PrimeSTAR GXL Premix, Clontech). The resulting DNA amplicon, along with a wildtype DNA amplicon, were sent to Quintara Biosciences for Sanger sequencing. The chromatographs of each sequencing run were used for TIDE (Tracking of Indels by Decomposition) analysis (https://tide.deskgen.com) and the cutting efficiency of hESCs nucleofected with FEV-KO gRNA was then determined. The FEV-KO hESC clonal line used in had a 1-bp deletion in one allele and a 1-bp insertion in the second allele, leading to a homozygous mutation in the *FEV* locus.

#### Human embryonic stem cell culture and differentiation to the beta cell lineage

FEV-KO and -WT hESC lines were maintained as clusters in suspension in mTeSR1 (STEMCELL Technologies) in 500 mL spinner flasks (Corning, VWR) on a magnetic stir plate (Dura-Mag) within a 37 °C incubator at 5% CO2, 100% humidity, and a rotation rate of 60 rpm. Cells were screened for mycoplasma contamination using the MycoProbe Mycoplasma Detection Kit (R&D Systems), according to the manufacturer’s instructions. Beta-like cells were generated as previously described (Millman et al., 2016; Pagliuca et al., 2014). In brief, single hESCs were seeded into a spinner flask at a density of 1e6 cells/mL in mTeSR1 media containing 10 μM Rock inhibitor Y-27632 (STEMCELL Technologies) to allow formation of clusters. Differentiation was initiated 72 h later and was achieved in a step-wise fashion using the following growth factors and/or small molecules: Stage 1!Day 1-3) medium : 500 mL MCDB 131 (Corning, 15-100-CV) + 0.22 g glucose (MilliporeSigma, G7528) + 1.23 g sodium bicarbonate (MilliporeSigma, S5761) + 10 g fatty-acid free bovine serum albumin (FAF-BSA) (Lampire Biological Laboratories, 7500812) + 10 μL Insulin-Transferrin-Selenium-Ethanolamine (ITS-X) (Invitrogen, 51500056) + 5 mL GlutaMAX (Invitrogen, 35050079) + 5 mL Penicillin-Streptomycin (P/S) solution (Corning, 30-002-CI). Stage 2 !Day 4-6) medium: 500 mL MCDB 131 + 0.22 g glucose + 0.615 g sodium bicarbonate + 10 g FAF-BSA + 10 μL ITS-X + 5 mL GlutaMAX + 0.022 g vitamin C (MilliporeSigma, A4544) + 5 mL P/S. Stage 3!Day 7-8) / 4!Day 9-13) medium: 500 mL MCDB 131 + 0.22 g glucose + 0.615 g sodium bicarbonate + 10 g FAF-BSA + 2.5 mL ITS-X + 5 mL GlutaMAX + 0.022 g vitamin C + 5 mL P/S. Stage 5!Day 14-20) medium:500 mL MCDB 131 + 1.8 g glucose + 0.877 g sodium bicarbonate + 10 g FAF-BSA + 2.5 mL ITS-X + 5 mL GlutaMAX + 0.022 mg vitamin C + 5 mL P/S + 5 mg heparin (MilliporeSigma, H3149). Stage 6 !Day 21-31) medium: 500 mL CMRL 1066 supplemented (Corning, 99-603-CV) + 10% Fetal Bovine Serum (FBS) (Corning, MT-35-011-CV) + 5 mL P/S.

#### Flow cytometric analysis of stem cell-derived cells

Stem cell-derived clusters at various stages of differentiation were washed in PBS (Corning, 21-040-CV) and dissociated with Accumax™ (Innovative Cell Technologies Inc, AM105) at 37 °C for the following times at each stage: 5 minutes (pluripotency), 5-7 minutes (End Stage 1), 7-9 minutes (End Stage 3), 9-11 minutes (End Stage 4), 11-13 minutes (End Stage 5), 12-15 minutes (Stage 6). Cells were then fixed with 4% PFA for 10 minutes at RT and spun down at 1,200 rpm for 5 minutes and resuspended in PBS. Cells were filtered through a 37 µm cell strainer (Corning, 352235) on ice, washed with 1X Permeabilization Buffer (00-8333-56, Invitrogen™) and spun down at 1200 rpm for 5 minutes at 4° C. Cells were stained with flow antibodies, then diluted in CAS Blocking Buffer (Invitrogen, 8120) containing 0.2% Triton-X, 5% NDS, and 1% bovine serum albumin (BSA) O/N at 4° C. The following morning, cells were washed with Permeabilization Buffer, spun down at 1,500 rpm for 5 minutes at 4° C, resuspended in FACS buffer (PBS, 1% FBS and 2mM EDTA), and analyzed on an LSR-II flow cytometer (BD Biosciences). At the end of each differentiation stage, FEV-KO or -WT hESC-derived cells were subjected to flow cytometric analyses, as described above. Data was analyzed with FlowJo software (Tree Star Inc.)

#### Quantitative RT-PCR

hESCs were collected from various stages of directed differentiation and subjected to RNA extraction using the RNeasy Mini Kit (QIAGEN 74106). Reverse transcription was performed with the Clontech RT-PCR kit. RT-PCR was run on a 7900HT Fast Real-Time PCR instrument (Applied Biosystems) with Taqman probes for *FEV* (assay ID: Hs00232733_m1) and *GAPDH* (assay ID: Hs02758991_g1) in triplicate. Expression of *FEV* was normalized to *GAPDH*.

#### Bulk-RNA sequencing

FEV-KO and -WT clusters were collected from four independent batches of stem cell differentiation at Stage 6 day 10 (S6D10). 2e6 cells were lysed in 350 ul RLT buffer (QIAGEN, 79216) and stored at -80 °C before RNA extraction. RNA was purified with RNeasy Mini Kit (QIAGEN, 74106). Samples with a RNA Integrity Number (RIN) greater than 9 were advanced to library construction using poly-A enrichment. Sequencing was conducted on a NovaSeq 6000 PE150 platform with the following parameters: Read 1 - 150 cycles, Index 1 i7 - 8 cycles, Index 2 i5 - 8 cycles, Read 2 - 150cycles. The resulting files were mapped to the reference genome (GRCh38) with STAR (v2.6.1d) and counts were generated with FeatureCounts (v1.26.0-p3). Differentially expressed genes were calculated with the DESeq2 (v1.26.0) workflow (Love et al., 2014).

### QUANTIFICATION AND STATISTICAL ANALYSIS

To analyze the population dynamics *in vivo* of each novel endocrine progenitor population over developmental time, nine samples of human fetal tissue from 8w (n = 3), 12w (n = 3), and 18w (n = 3) were stained using multiplexed *in situ* hybridization and immunofluorescence. For each biological sample, images from five areas were taken at random and processed with the maximum intensity z-projection function with the ImageJ software package. Adjustments to brightness and contrast were applied equally across images in a given series. The number of cells corresponding to each cell state was manually counted in each biological sample from the five image areas using the Image J plug-in Cell Counter. The proportion of each cell state present was then calculated using the sum of cells corresponding to all cell states as the denominator, and cells that scored positive for a given cell state as the numerator. Data were presented as Mean ± SEM (n = 3). Graphs were generated in GraphPad software (Prism 8). When assessing the proportional changes of *FEV*+*NEUROG3*+ progenitor cells over developmental time, an unpaired t-test was used to determine the statistical significance of the difference between the ratio of *FEV*+*NEUROG3*+ progenitor cells at 8w and at 12w, as well as the ratio difference between 12w and 18w.

To assess differences between stem cell-derived cells from FEV-KO vs. -WT hESCs, data were quantified from flow cytometric analyses. At the early beta-like cell stage (Stage 6, day 4), the proportion of C-PEP+/NKX6-1+ double positive cells in FEV-KO or -WT cells was analyzed from three independent batches of differentiation. Statistical significance of difference between the two groups was determined using the paired t-test in GraphPad software (Prism 8). Data were presented as Mean ± SEM. To quantify the proportional changes of hormone-producing cells upon *FEV* knockout, immunofluorescence images of FEV-KO and -WT clusters from two independent batches of differentiation were manually counted using ImageJ software, and differences in cell proportion between WT and FEV-KO were assessed with an unpaired t-test in GraphPad software (Prism 8).

## Acknowledgments

This work was supported by grants to J.B.S. from the NIH (R01DK118421) and JDRF (2-SRA-2019-773-S-B). S.M.D. was supported by the Kraft Family Fellowship to the UCSF Diabetes Center, the UCSF Discovery Fellows Program, NIH NIGMS IMSD Grant #R25GM056847-23, and NIH/NIDDK diversity supplement R01DK118421-02S1. Z.L. was supported by the Jeffrey G. Klein Family Diabetes Fellowship to the UCSF Diabetes Center, and D.M.W. was supported by a National Science Foundation (NSF) Graduate Research Fellowship and the Diabetes, Endocrinology, and Metabolism Training Program from the NIH (T32 DK007418). E.C. was supported by a fellowship from the California Institute of Regenerative Medicine (CIRM) (SFSU EDUC2-08391). A.L.G. is a Wellcome Senior Fellow in Basic Biomedical Science. A.L.G. is funded by the Wellcome (200837) and National Institute of Diabetes and Digestive and Kidney Diseases (NIDDK) (U01-DK105535; U01-DK085545, UM1DK126185) and the Stanford Diabetes Research Center (NIDDK award P30DK116074).

The authors would like to acknowledge expert technical assistance from the UCSF Parnassus Flow Cytometry Core, the UCSF Broad Imaging Core, the UCSF Institute for Human Genetics Core, and Gregory Szot of the UCSF Islet Production Core. We thank members of the Sneddon Laboratory, Dr. Alison May, Dr. Nicole Krentz and Steven Cincotta for thoughtful suggestions.

## Author Contributions

Z.L., S.M.D., and J.B.S. conceived the study. Z.L. and D.M.W. performed sample preparation and cell enrichment for scRNA-Seq. Z.L. performed nuclei extraction for snATAC-Seq. S.M.D and Z.L. performed computational analysis. S.M.D., Z.L. and J.B.S interpreted computational results. K.S.Y. and A.D.T. provided access to, and assistance with, CellFindR software. N.E. and Y.S. assisted with snATAC Seq analysis and interpretation. A.L.G. and H.S. performed computational studies on the identification of development-specific type 2 diabetes GWAS risk loci. Z.L. and E.C. performed and analyzed *in situ* hybridization and immunofluorescence of human fetal tissue. D.M.W., G.P., and J.O.B. generated CRISPR/Cas9 edited FEV-KO and WT cell lines; S.A.R., S.M.D., Z.L, and E.C. directed their differentiation into beta-like cells. Z.L., E.C., and S.A.R. performed flow cytometric analyses. S.M.D. and Z.L assembled figures. S.M.D., Z.L., A.L.G., and J.B.S. wrote the manuscript. S.M.D., Z.L., S.A.R., E.C. and H.S. contributed methods. All authors approved the content and submission of the manuscript. J.B.S. oversaw the project, supervised all personnel, and acquired funding.

## FIGURE/FIGURE LEGENDS

**Figure S1:**
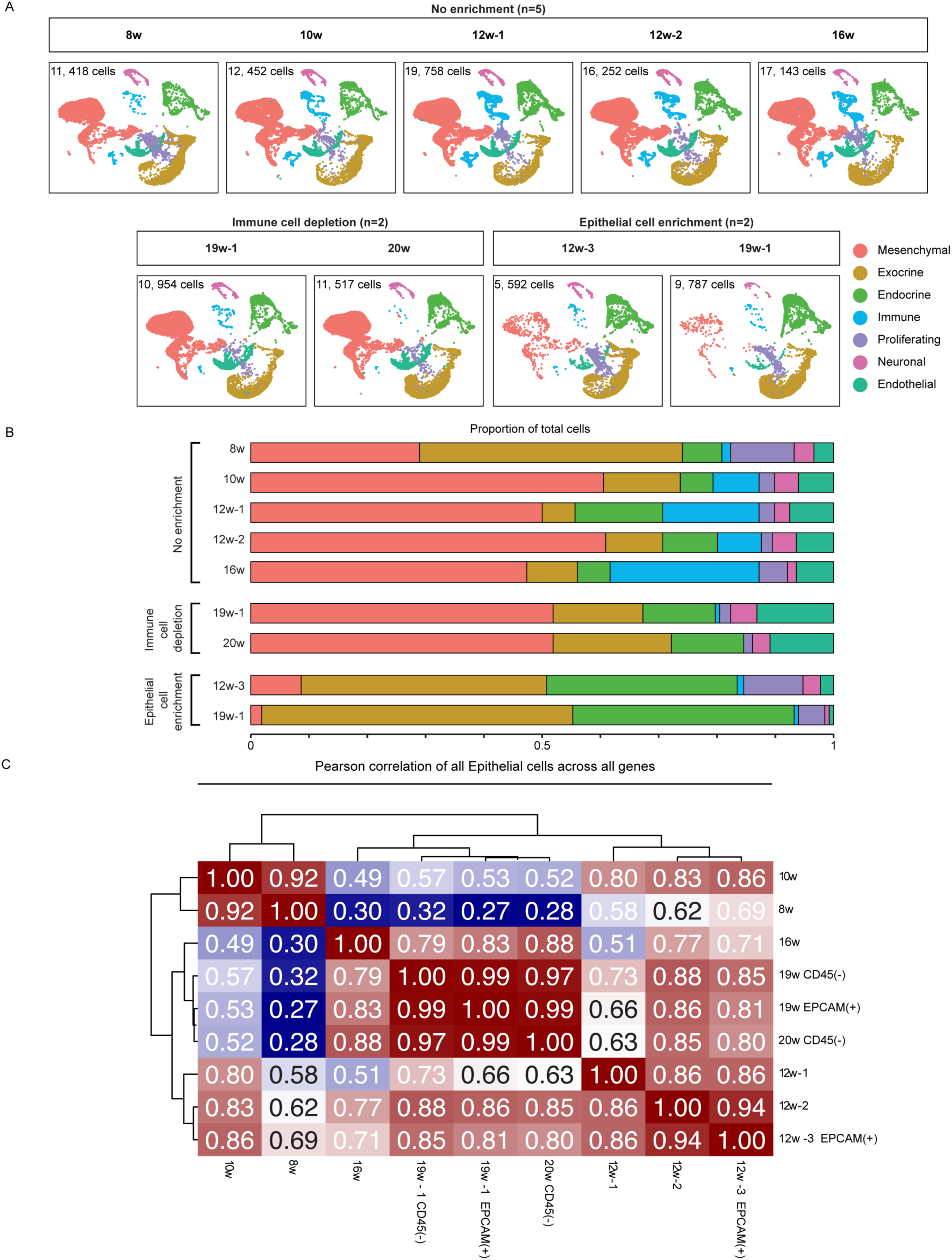
CellFindR clustering and Broad Group proportional representation in the human fetal pancreas, parsed by individual sample. (A) Split UMAP visualization shows contribution of each sample to the overall merged dataset, with cells colored according to their Broad Group identity. The number of cells contributing to the merged data set is listed on each UMAP. (B) Bar graph depicting the proportional representation of each Broad Group in each of the individual scRNA-Seq samples. (C) Heatmap depicting Pearson correlation of all Epithelial cells (Exocrine and Endocrine Broad Groups) across fetal scRNA Seq samples.

**Figure S2:**
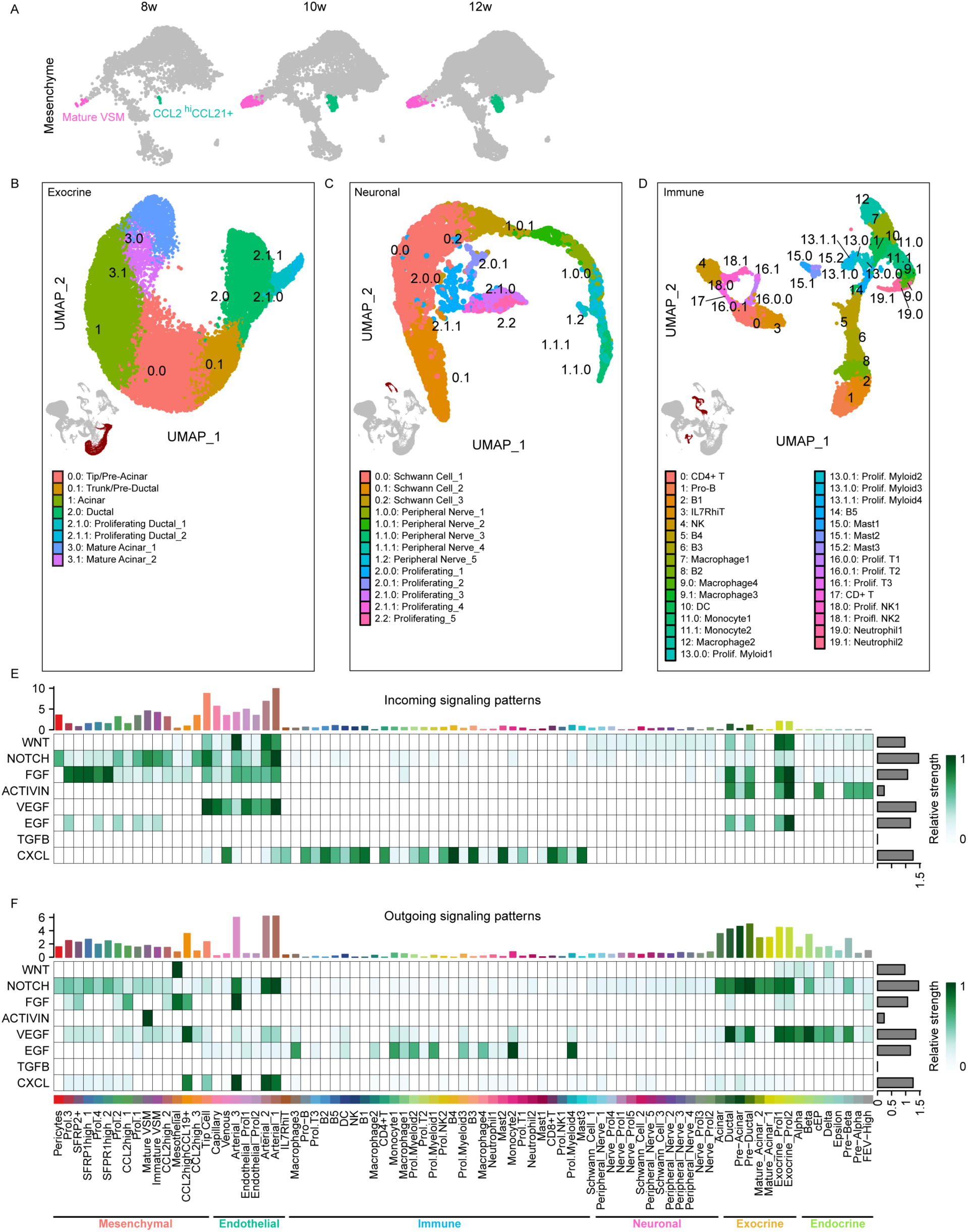
Novel populations discovered within the exocrine, neuronal, and immune compartments, and predicted intercellular communication between all Broad Group cell subtypes. (A)-(C) UMAP visualization of cell populations identified by CellFindR within the (A) exocrine, (B) neuronal, and (C) immune broad groups within the human fetal pancreas. (D)-(E) Predicted paracrine signaling pathways that mediate cell-cell communications between different Broad Groups, as inferred by CellChat analysis. (D) Plot of incoming signaling patterns depicts the predicted cell source of each paracrine ligand identified as significant. (E) Plot of outgoing signaling patterns plot reveals cell specificity of paracrine signaling receptors predicted to be significant. Bar graphs at the top of (D) and (E) represent the aggregate signaling for each cluster across all signaling pathways; bar graphs on the right represent the aggregate signaling strength of each signaling pathway across all clusters.

**Figure S3:**
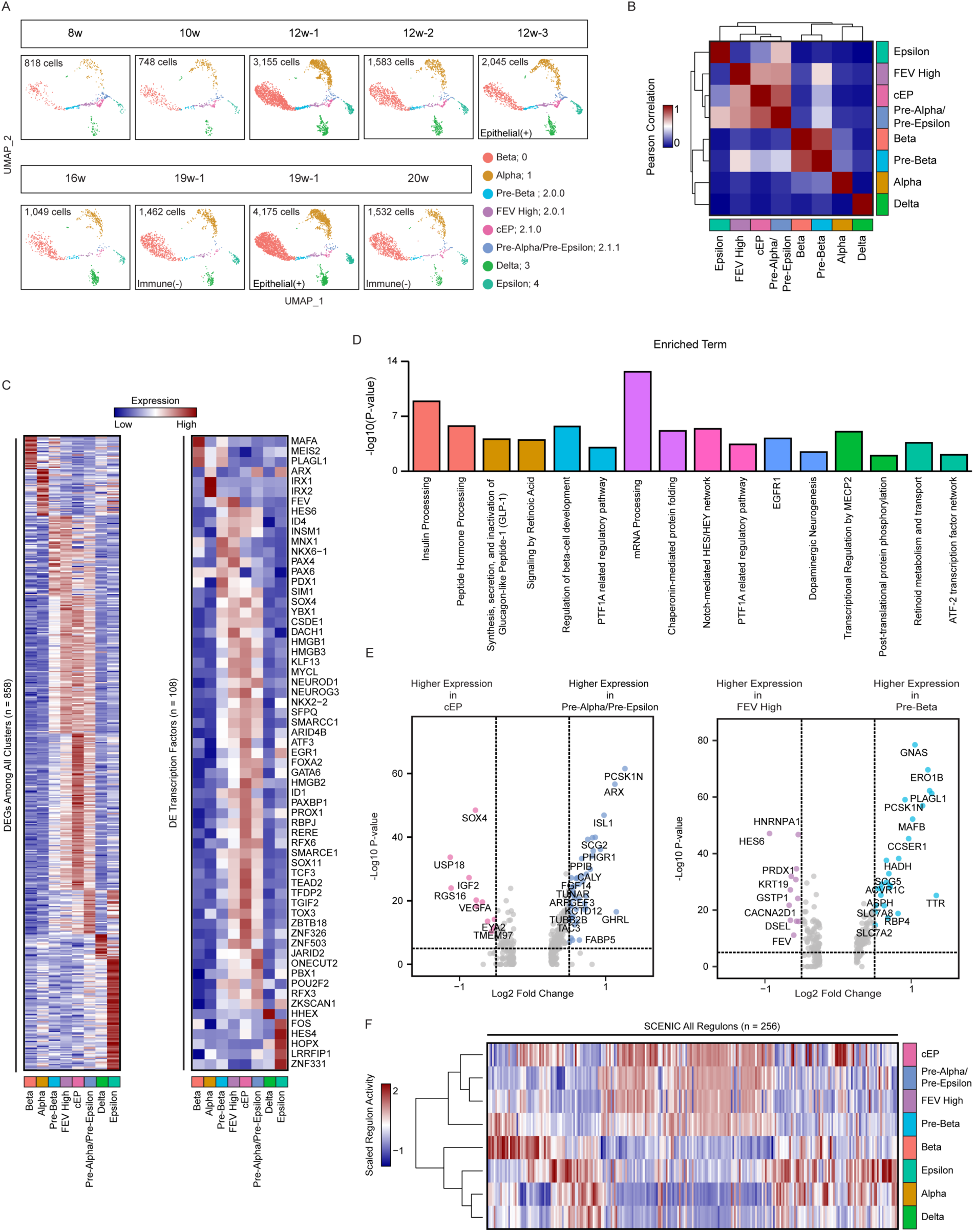
Transcriptomic and population analysis of human fetal endocrine cells. (A) Split UMAP visualization shows the contribution of each individual sample to the merged endocrine scRNA-seq dataset, with the number of cells listed and each cell colored according to its endocrine cluster identity. (B) Heatmap depicting the Pearson correlation between pairs of cell populations, based on the top 1,000 most highly variable genes among the endocrine clusters. (C) Heatmaps depicting expression levels of all differentially expressed genes among endocrine cells of the human fetal pancreas (left). All differentially expressed transcription factors are depicted in the heatmap on the right. (D) Pathway analysis of all genes with a log2-fold change in expression of at least 0.5 between a single endocrine cell population and all other endocrine cell populations of the human fetal pancreas. For each significant pathway, the clusters in which that pathway is active is colored to match the color in panel (A). (E) Volcano plots depicting pairwise comparisons of gene expression in Common Endocrine Progenitor (cEP) vs. Pre-Alpha/Pre-Epsilon cells (left panel) and Pre-Beta vs. FEV High cells (right panel). Genes with a log2-fold change of at least 0.5 are highlighted. (F) Heatmap depicting the regulon activity score of all 256 regulons identified as significant by SCENIC analysis. Clusters in (F) are annotated by their cluster color as shown in (A).

**Figure S4:**
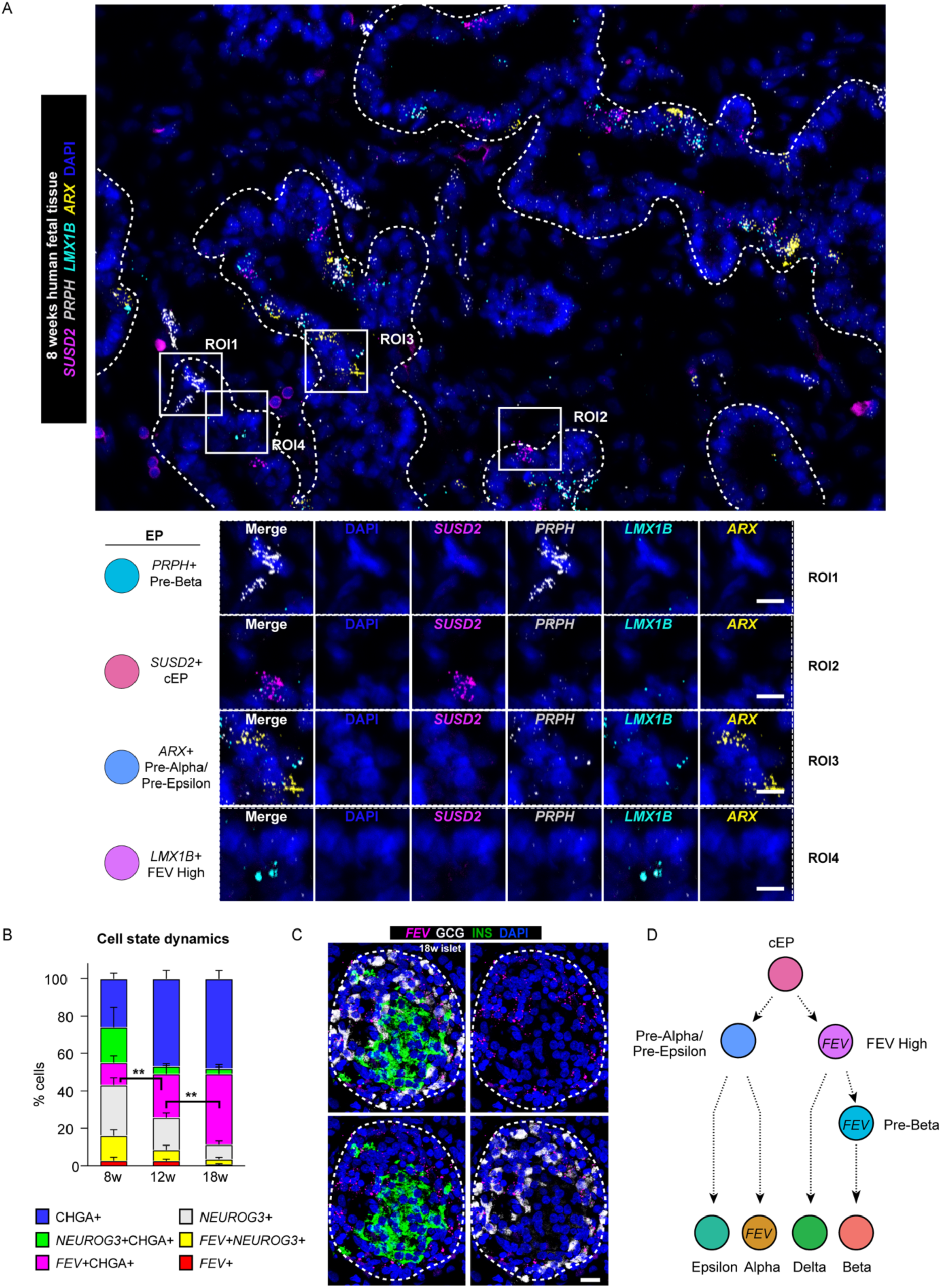
Confirmation *in vivo* of novel putative endocrine progenitor populations in the developing human pancreas. (A) Multiplexed staining of 8 w human fetal tissue with four probes against novel endocrine progenitor markers: *SUSD2* (red), *PRPH* (grey), *LMX1B* (cyan), and *ARX* (yellow); nuclei were counterstained with DAPI (blue). Regions of interest (ROIs) 1, 2, 3, and 4 highlight cells expressing single marker genes distinguishing Pre-Beta EP, cEP, Pre-Alpha/Pre-Epsilon EP, and FEV High EP cells, respectively. Dashed lines indicate epithelial border in the tissue. Scale bars, 10 um. (B) Dynamics of cell states assessed by staining for *FEV/NEUROG3*/CHGA, across developmental time. The relative proportion of *FEV*+CHGA+ cells significantly increased as development progressed. **, P-value < 0.01; graphs are presented as mean ± SEM (n=3 biological samples); significance was calculated using an unpaired t-test. (C) Multiplexed ISH/IF staining for *FEV* (magenta), INS (green), and GCG (grey) on 18 w tissue. An islet is circled with a dashed line. Scale bar, 25 um. (D) Diagram summarizing expression of *FEV* in endocrine progenitors and GCG+ alpha cells.

**Figure S5.**
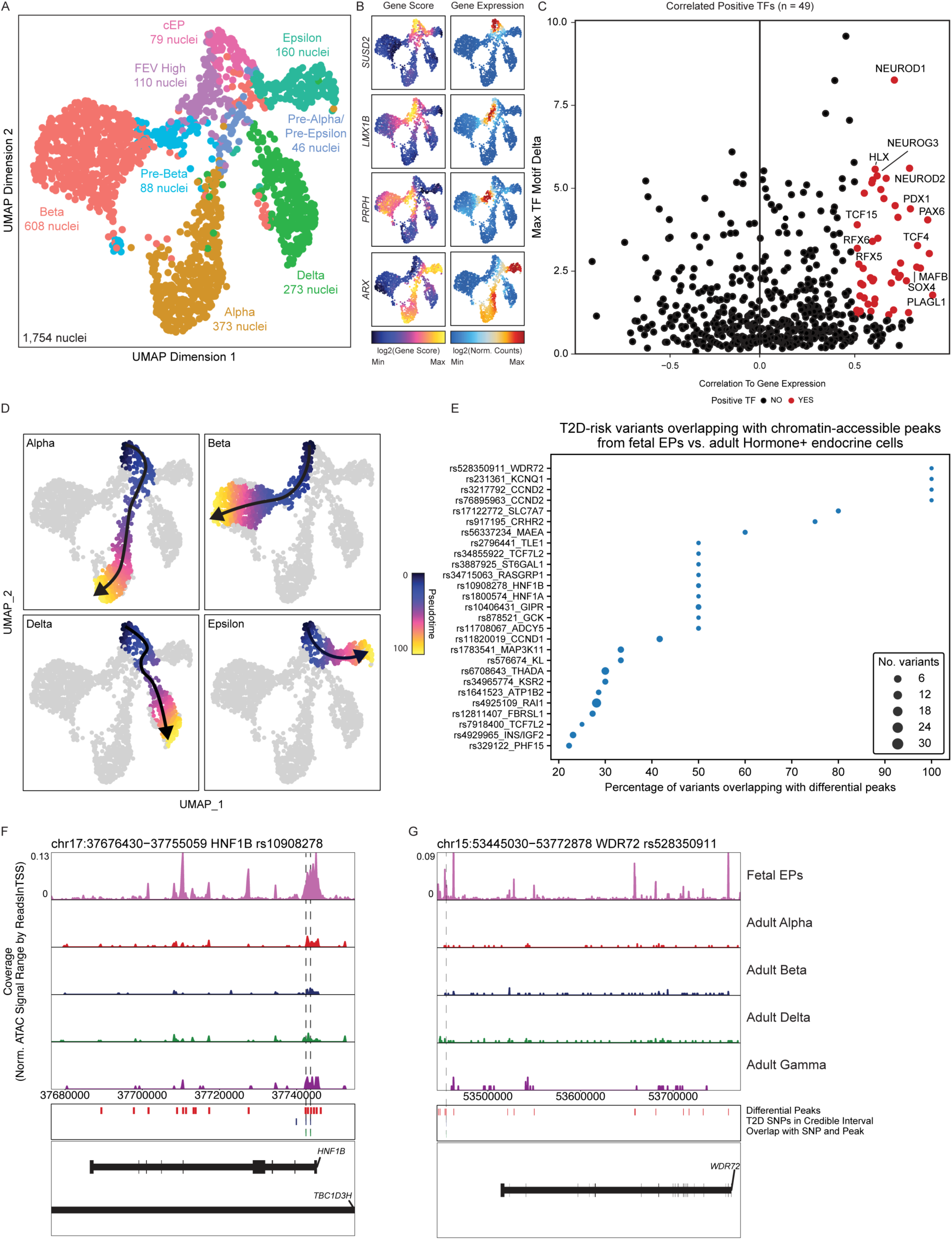
Identification of “positive” transcription factors across endocrine lineages using multi-omic analysis. (A) UMAP displaying snATAC-Seq endocrine cells annotated via unconstrained integration with the scRNA-Seq dataset from Figure 2. In contrast to Figure 5B, here the EP subpopulations are individually labeled and not pooled. (B) ATAC gene scores (left) and corresponding RNA expression values (right) of endocrine progenitor markers as defined in Figure 2C. (C) Plot depicting TFs scored as “positive” in all endocrine cells by correlation of chromVAR deviation and gene expression (n = 49 genes). (D) Lineage reconstruction based on snATAC-Seq data. (E) T2D-risk loci enriched in differentially accessible peaks of fetal EP cells as compared to all adult hormone+ cells. (F), (G) Track plots displaying accessibility of the *HNF1B* and *WDR72* loci among fetal EPs (top) vs. adult hormone-expressing cells.

**Figure S6:**
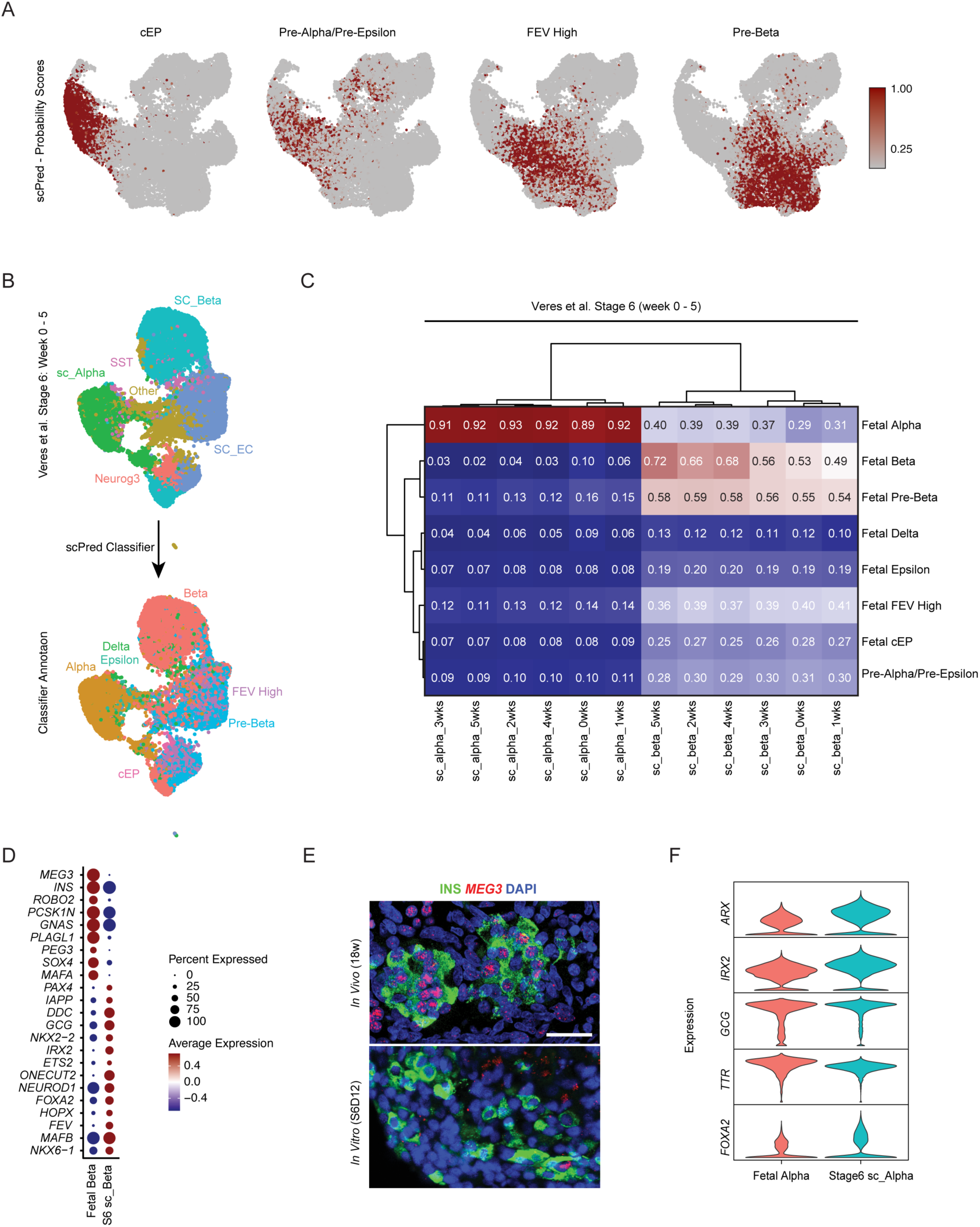
Transcriptional comparison of *in vitro* stem cell-derived vs. fetal endocrine cells. (A) Feature plots of *in vitro* stem cell-derived Stage 5 cells (Figure 6B) depicting scPred probability scores for the cEP, Pre-Alpha/Pre-Epsilon, FEV High and Pre-Beta human fetal clusters. (B) UMAP of cells generated at Stage 6 of the stem cell differentiation protocol, with annotation by Veres *et al*. (top) or by cell-based classifier scPred (bottom) based on cell type similarity to human fetal endocrine cells. (C) Heatmap depicting the Pearson correlation between cells generated at Stage 6 *in vitro* and fetal endocrine cell types, based on all shared genes. Stage 6 cell types are named according to the duration (number of weeks) that they were cultured in Stage 6. (D) Dot plot depicting differentially expressed genes between fetal beta cells and Stage 6 stem cell-derived beta-like cells (Fetal Beta and S6 sc_Beta, respectively). (E) Dual *in situ* hybridization/immunofluorescence staining for *MEG3* mRNA and INS protein, with DAPI staining nuclei, in 18 w human fetal tissue (top) and stem cell-derived clusters (Stage 6 day 12) (bottom). Scale bar, 25um. (F) Violin plots depicting the expression of known regulators of alpha cells in fetal and *in vitro* stem cell-derived Stage 6 alpha-like cells.

**Figure S7:**
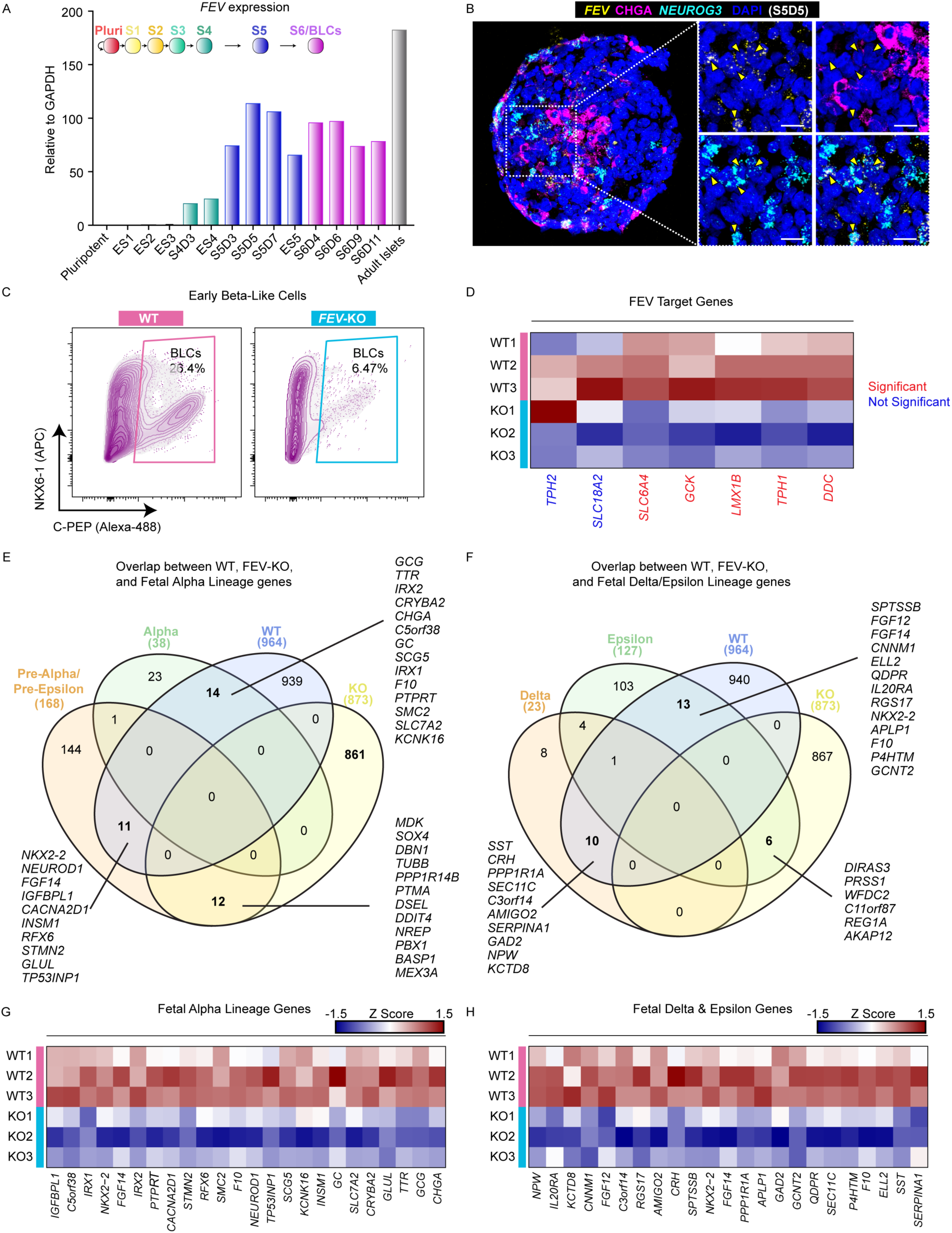
*FEV* marks endocrine progenitor cells *in vitro*, and its loss results in impaired endocrine development. (A) Beta-like cells (BLCs) were generated from pluripotent (Pluri) human embryonic stem cells (hESCs) using a six stage differentiation protocol. *FEV* mRNA expression was measured throughout the differentiation by qRT-PCR Taqman analysis. Adult human islets are included as control. (B) Dual *in situ* hybridization (ISH) and immunofluorescence (IF) staining of a cluster of *in vitro* hESC-derived endocrine stage cells (Stage 5, day 5) cells to detect *FEV* mRNA (yellow) and *NEUROG3* mRNA (cyan) along with CHGA protein (magenta). Nuclei were counterstained with DAPI (blue). Scale bar: 25um. Yellow arrowheads indicate *FEV+/NEUROG3+* double-positive putative endocrine progenitors. (C) Representative flow cytometry analysis for beta cell lineage markers NKX6-1 and C-PEPTIDE (C-PEP) at the early BLC stage (Stage 6, day 4). (D) Heat map showing gene expression levels of known targets of *FEV* in FEV-KO and WT cells at the BLC stage (Stage 6, day 10). (E) Four-way Venn diagram depicting the overlap among DEGs (defined as log2 FC > 0.5) distinguishing fetal alpha lineage cell populations and those identified from a comparison of in WT vs. FEV-KO BLCs. (F) Four-way Venn diagram depicting the overlap among DEGs distinguishing fetal delta or epsilon cell populations and those distinguishing WT vs. KO BLCs. (G) Heatmap depicting the expression levels of fetal alpha lineage genes and (H) the expression levels of fetal delta and epsilon lineage genes in paired differentiations of WT and KO cells at the BLC stage (Stage 6, day 10).

